# An adapted stem cell-derived microglia protocol for the study of microgliopathies and other neurological disorders

**DOI:** 10.1101/2023.09.08.556933

**Authors:** Marie-France Dorion, Diana Casas, Moein Yaqubi, Peter Fleming, Carol X.-Q. Chen, Valerio E. C. Piscopo, Michael Nicouleau, Taylor M. Goldsmith, Irina Shlaifer, Adam MacDonald, Roy W. R. Dudley, Jeffrey A. Hall, Jack P. Antel, Jo Anne Stratton, Thomas M. Durcan, Roberta La Piana, Luke M. Healy

## Abstract

**Background:** Adult-onset leukoencephalopathy with axonal spheroids and pigmented glia (ALSP) is a primary microgliopathy caused by pathogenic variants in the colony-stimulating factor 1 receptor (*CSF1R*) gene. Since CSF1R signaling is crucial for microglia development, survival and function, induced pluripotent stem cell-derived microglia (iMGL) represent an excellent tool in studying microglial defects caused by ALSP patient-specific *CSF1R* variants.

**Methods:** Serial modifications to an existing iMGL protocol were made, including but not limited to changes in growth factor combination to drive microglial differentiation, until successful derivation of microglia-like cells from an ALSP patient carrying a c.2350G > A (p.V784M) *CSF1R* variant. Using healthy control lines, the quality of the new iMGL protocol was validated through cell yield assessment, measurement of microglia marker expression, transcriptomic comparison to primary microglia, and evaluation of inflammatory and phagocytic activities. Similarly, molecular and functional characterization of the ALSP patient-derived iMGL was carried out in comparison to healthy control iMGL.

**Results:** The newly devised protocol allowed the generation of iMGL with enhanced transcriptomic similarity to primary human microglia and with higher phagocytic and inflammatory competence at ∼3-fold greater yield compared to the original protocol. Using this protocol, decreased CSF1R autophosphorylation and cell surface expression was observed in iMGL derived from the ALSP patient compared to those derived from healthy controls. Additionally, ALSP patient-derived iMGL presented a migratory defect accompanying a temporal reduction in purinergic receptor P2Y12 (*P2RY12*) expression. Finally, ALSP patient-derived cells showed surprisingly high phagocytic capacity, which was associated with higher lysosomal content.

**Conclusions:** We optimized a pre-existing iMGL protocol, generating a powerful tool to study microglial involvement in human neurological diseases. Using the optimized protocol, we have generated for the first time iMGL from an ALSP patient carrying a pathogenic *CSF1R* variant, with preliminary characterization pointing toward functional alterations in migratory and phagocytic activities.

## Background

Microglia are resident myeloid cells of the central nervous system that originate from yolk sac progenitor cells. Following brain colonization, microglia aid in the development of neuronal networks and myelination. As mononuclear phagocytes, they have crucial functions in the removal of excess neurons and synapses during development, as well as in the clearance of dying cells and debris. These highly motile cells express a diverse range of immune sensing receptors, microglia mediate the immune surveillance of brain parenchyma (1).

Recent advances in the human microglia field have been driven by the development of several induced pluripotent stem cell-derived microglia (iMGL) protocols starting in 2017 (2). Derivation of microglia from induced pluripotent stem cells (iPSCs) is generally carried out in two steps: 1) production of hematopoietic precursor cells, and 2) microglial differentiation. In particular, the iMGL protocol developed by Abud *et al.* (3) and later simplified by McQuade *et al.* (hereafter referred to as version “2.0” (4)) relies on the use of macrophage colony-stimulating factor (M-CSF), interleukin-34 (IL-34) and transforming growth factor-beta 1 (TGF-β1) for microglial differentiation. These growth factors are important for the acquisition of a unique molecular identity that distinguishes microglia from other myeloid populations (5–7).

M-CSF and IL-34 are endogenous ligands of the receptor tyrosine kinase colony-stimulating factor 1 receptor (CSF1R) essential for both the development and the maintenance of microglia pool in the brain through self-renewal (7–11). In mice, *Csf1r* knockout results in an almost complete failure of microglia development (12). In human, mono-allelic pathogenic variants in the *CSF1R* gene have been associated with adult-onset leukoencephalopathy with axonal spheroids and pigmented glia (ALSP). This rare autosomal dominant disease is pathologically characterized by vacuolating and demyelinated white matter especially in the frontal regions and corpus callosum, axonal degeneration and swelling (spheroids) and pigmented myeloid cells in the brain. Affected individuals present with psychiatric, cognitive and motor symptoms usually in the 4^th^ decade, and rapidly deteriorate to death on average 7 years after disease onset (13, 14). The impact of *CSF1R* pathogenic variants on microglia function and brain abnormalities remains poorly understood, and no treatment exists to date.

As most iMGL protocols including the 2.0 protocol developed by McQuade *et al.* heavily rely on CSF1R signaling for successful microglia differentiation (4) and survival (15), it did not appear possible to actively study *CSF1R*-mutated microglia using such protocols. We made serial modifications to the microglia differentiation medium used in the 2.0 protocol to ensure successful derivation of microglia-like cells from an ALSP patient harboring a pathogenic c.2350G > A *CSF1R* variant. This protocol, which conserved the simplicity of the 2.0 protocol all the while improving the overall functional competence of the resulting cells, will be referred to as the “2.9” protocol since it was the 9^th^ iteration of the 2.0 protocol.

Herein, we present the new 2.9 protocol and the characterization of the resulting cells (“iMGL 2.9”) in comparison to cells generated using the original protocol (“iMGL 2.0”) and primary human microglia. This new protocol was used to carry out the cellular and molecular phenotyping of ALSP patient-derived iMGL carrying a heterozygous pathogenic variant in the *CSF1R* gene, which we failed to achieve using the 2.0 protocol due to a failure in differentiation.

## Methods

### IPSC lines

Seven control iPSC lines were used in this study. Characteristics of each line are presented in Additional file 2: Table S1. Generation of iPSC lines was done following McGill University Health Centre’s ethical guidelines (project# 2019-5374) with written consent from donors. Cells were maintained in mTeSR^TM^ Plus (STEMCELL Technologies) or in Essential 8^TM^ media on Corning^TM^ Matrigel^TM^ hESC-Qualified Matrix -coated dishes, with subculturing every five to seven days using standard protocols (16).

For the generation of iPSCs from a *CSF1R*-mutated ALSP patient, peripheral blood mononuclear cells (PBMCs) were collected through the intermediary of C-BIG Repository, Montreal, Canada, with written consent from the donor and following McGill University Health Centre’s ethical guidelines. PBMCs were reprogrammed into iPSCs using previously established episomal reprogramming method (16). Briefly, PBMCs were nucleofected with episomal plasmids (pEV-OCT4-2A-SOX2, pEV-MYC, pEV-KLF4, and pEV-BC-XL) and resulting iPSC colonies were selected based on the acquisition of stem cell-like morphology for expansion and cryopreservation. Genomic integrity was then verified by karyotyping and quantitative polymerase chain reaction (qPCR)-based assays as previously described (16).

### Generation of iMGL

#### Hematopoietic differentiation

Differentiation of iPSCs into iPSC-derived hematopoietic progenitor cells (iHPCs) was carried out using STEMdiff^TM^ Hematopoietic kit (STEMCELL Technologies) as previously described (4) with minor modifications. On day -1, iPSCs were detached using Gentle Cell Dissociation Reagent (STEMCELL Technologies) and plated in Matrigel^TM^-coated 6-well plates, in mTeSR^TM^ Plus or in Essential 8^TM^ media supplemented with Y27632 (10 μM, Selleckchem). Several seeding densities should be tested for every differentiation batch, aiming in the range of 1-5 small colonies per cm^2^ at the start of the differentiation. On day 0, media were replaced with STEMdiff^TM^ hematopoietic medium A (2 mL/well). On day 2, half the volume of the cell supernatants (1 mL/well) was replaced with fresh medium A. On day 3, media was fully replaced with STEMdiff^TM^ hematopoietic medium B (2 mL/well). On day 5 and 7, half the volume of the cell supernatants (1 mL/well) was replaced with fresh medium B. On day 9, 1 mL/well of fresh medium B was added. On day 10, cell supernatants containing the floating iHPCs were collected and spun down at 300 g for 5 minutes. 1 mL/well of cell-free conditioned media, along with 1 mL/well of fresh medium B, were put back on the iHPC culture. The collected pellet of iHPCs was processed either for cryopreservation (using Bambanker, Fujifilm Wako Chemicals) or for microglial differentiation. iHPCs were similarly harvested on day 12.

#### Microglial differentiation

On day 0, iHPCs were resuspended at a density of 50 000-100 000 cells/mL in microglia differentiation medium 2.0 or 2.9 (Table 1) and plated on Matrigel^TM^-coated 6-well plates (2 mL/well). From day 0 to day 10, the culture was supplemented with 1 mL/well of differentiation media every other day. On day 12, 6 mL/well of cell supernatants were collected and spun down at 300 g for 5 minutes. The collected cells were resuspended in 1 mL/well of differentiation media and placed back in culture. From day 12 to day 22, the culture was supplemented with 1 mL/well of differentiation media every other day. On day 24, 6 mL/well of cell supernatants were collected and spun down at 300 g for 5 minutes. The collected cells were resuspended in 1 mL/well of differentiation media and put back in culture. Cells were considered mature on day 28, and were maintained in a similar manner with media supplementation every other day until needed. Throughout the protocol, cells were maintained at 37 °C under a 5% CO2 atmosphere. For any downstream experiments, cells were detached using PBS with 2 mM ethylenediaminetetraacetic acid (EDTA; 10-minute incubation) and replated at a density of 7.5 × 10^5^ cells per cm^2^.

**Table 1.**
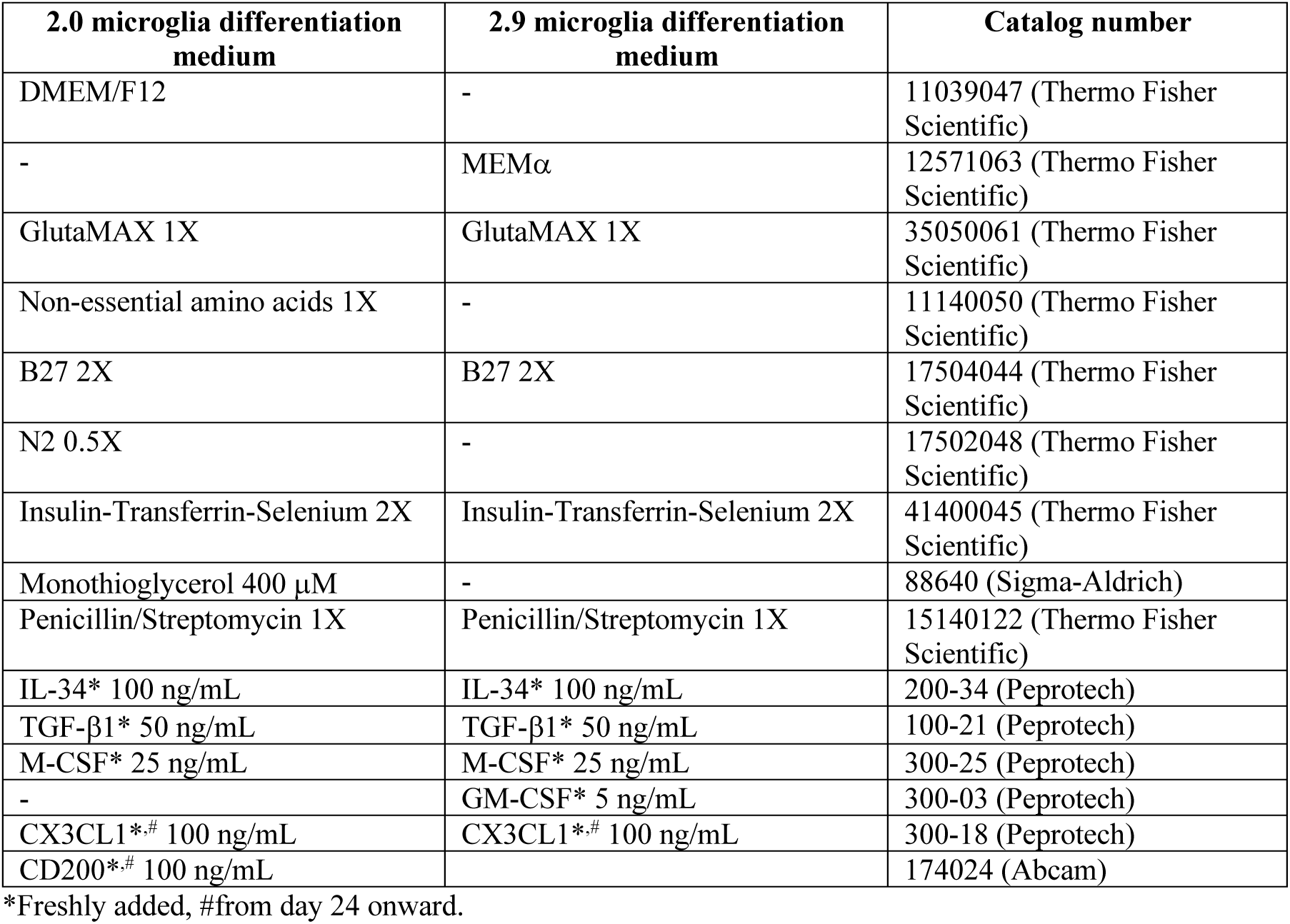
Composition of the 2.0 and 2.9 microglia differentiation media.

### Primary microglia isolation and culture

Human brain tissues from 4-to 65-year-old female and male epilepsy patients were obtained from the Montreal Neurological Institute, Montreal, Canada (adult donors) and the Montreal Children’s Hospital, Montreal, Canada (pediatric donors), with written consent and under local ethic boards’ approval. Tissues were from sites distant to the suspected primary epileptic foci. Isolation of glial cells was carried out as previously described (17) through mechanical and chemical digestion, followed by Percoll^®^ (Sigma-Aldrich) gradient centrifugation. Microglia were further purified by taking advantage of the differential adhesive properties of the glial cells. Cells were maintained in Minimum Essential Medium (MEM; Sigma-Aldrich) supplemented with 5% fetal bovine serum (FBS; Wisent Bio Products), 1% P/S, 0.1% glucose and 1X GlutaMAX^TM^. Fetal microglia were isolated similarly (17) from second trimester fetal brain tissues obtained from Centre Hospitalier Universitaire Sainte-Justine, Montreal, Canada, with maternal written consent and under local ethic boards’ approval. Fetal microglia were cultured in Dulbecco’s Modified Eagle’s Medium (DMEM; Thermo Fisher Scientific) supplemented with 5% FBS, 1% P/S and 1X GlutaMAX^TM^.

### Peripheral blood mononuclear cell-derived macrophages (PBMC-Mχπ)

PBMCs were isolated from whole blood by Ficoll gradient centrifugation. Monocytes were then collected through magnetic activated bead sorting of cluster of differentiation 11b (CD11b) -positive cells, and cultured at a density of 5-10 × 10^5^ cells/cm^2^ (day 0) in RPMI-1640 (Thermo Fisher Scientific) with 10% FBS, 1% P/S, 1X GlutaMAX^TM^ and 30 ng/mL M-CSF (Peprotech). Cells were matured for 8 days, with media supplementation on day 4 and day 7.

### Immortalized cell lines

hTERT RPE-1 and THP-1 cells were from American Type Cell Collection. hTERT RPE-1 cells were maintained in DMEM/F12 with 10% FBS and 1% P/S with subculturing every four days. THP-1 cells were maintained in RPMI-1640 with 10% FBS, 1% P/S and 1X GlutaMAX^TM^ with subculturing every five days.

### Adhesion assay

A crystal violet kit (Abcam) was used to assess cell adhesion following the manufacturer’s protocol. Cells were first seeded on a 96-well plate. After 48 hours, cells were washed with washing buffer to remove non-adherent cells, prior to incubation with crystal violet. After washing off the excess crystal violet, cells were incubated in the solubilization buffer. Absorbance was measured at 560 nm on a SpectraMax® iD3 microplate reader (Molecular Devices).

### Immunocytochemistry

Cells were fixed in 4% paraformaldehyde and permeabilized/blocked using PBS with 3% goat or donkey serum and 0.2% triton X-100 (Sigma Aldrich). Cells were incubated at 4°C overnight with primary antibodies against the following targets: ionized calcium binding adaptor molecule 1 (IBA1; #NC9288364, Fujifilm Wako Chemicals at 1:1000), PU.1 (#2258, Cell Signaling at 1:250), Nanog (#ab21624, Abcam at 1:500), Tra-1-60 (#60064, STEMCell Technologies at 1:200), stage-specific embryonic antigen-4 (SSEA-4; #sc-21704, Santa Cruz Biotechnologies at 1:200), octamer-binding transcription factor 3/4 (OCT3/4; #sc-8628, Santa Cruz Biotechnologies at 1:500), lysosomal-associated membrane protein 1 (LAMP1; #9091, Cell Signaling at 1:200) or cluster of differentiation 68 (CD68; #M0814, Dako Omnis at 1:200). Cells were then incubated with secondary antibodies and 1 μg/mL Hoechst 33342 for one hour. The proportion of cells with positive stain was determined using a CellInsight CX5 High Content Screening Platform (Thermo Fisher Scientific). All conditions were assessed in triplicate.

### Ribonucleic acid (RNA) -sequencing

TRIzol (Thermo Fisher Scientific) was used to extract RNA, followed by cleaning using a RNeasy mini kit (Qiagen). Quality control of the RNA samples, as well as the library preparation by poly(A) enrichment and RNA-sequencing were performed by Genome Quebec, Montreal, Canada. RNA-sequencing was performed using an Illumina NovaSeq 6000 with a read depth of 50 million reads per sample. Canadian Center for Computational Genomic’s pipeline GenPipes (18) was used to align the raw files and quantify the read counts. Briefly, raw fastq files were aligned to the GRCh38 genome reference using STAR aligner (19) with default parameters and raw reads were quantified using HTseq count (20). Differential expression gene (DEG) analysis was carried out using the DESeq2 package (21). DEGs were identified using an adjusted p-value cutoff of 0.05. Gene ontology (GO) enrichment analyses were performed using PANTHER overrepresentation test, on the web-based tool offered by the Gene Ontology Consortium. Principal component analysis (PCA) was carried out using the Python module sklearn and visualized using Matplotlib. Histograms were generated using GraphPad Prism 9.0 software.

### Quantitative reverse transcription polymerase chain reaction (qRT-PCR)

Following RNA extraction, reverse transcription was performed using Moloney murine leukemia virus reverse transcriptase (Thermo Fisher Scientific). Real-time PCR was performed using TaqMan assays (Thermo Fisher Scientific) on a QuantStudio^TM^ 5 real-time PCR system (Thermo Fisher Scientific). The 2^-ýCt^ method was used to analyze the data using glyceraldehyde 3-phosphate dehydrogenase (*GAPDH*) and tyrosine 3-monooxygenase/tryptophan 5-monooxygenase activation protein zeta (*YWHAZ*) as controls.

### Flow cytometry

Cells were blocked with Human TrueStain FcX and TrueStain Monocyte Blocker (Biolegend) and stained with the following antibodies: anti-cluster of differentiation 34 (CD34; clone #561, Biolegend), anti-cluster of differentiation 43 (CD43; clone #CD43-10G7, Biolegend), anti-cluster of differentiation 14 (CD14; clone #HCD14, Biolegend), anti-CSF1R (clone #61708, R&D Systems), anti-CX-3-C motif chemokine receptor 1 (CX3CR1; clone #2A9-1, Biolegend), anti-Mer tyrosine kinase (MERTK; clone #125518, R&D Systems), anti-purinergic receptor P2Y12 (P2RY12; clone #S16001E, Biolegend) or anti-toll-like receptor 4 (TLR4; clone #610029, R&D Systems). All antibodies were titrated using negative control cells that don’t or poorly express the target protein. Appropriate forward and side scatter profiles were used to exclude debris and doublets from the analysis. Dead cells were excluded based on LIVE/DEAD^TM^ Fixable Aqua (Thermo Fisher Scientific) staining. Readings were done on an Attune^TM^ Nxt Flow Cytometer and analyzed/visualized using FlowJo^TM^ software.

### Phagocytosis assay

Human α-synuclein preformed fibrils (22), myelin debris (23) and immunoglobulin G-opsonized red blood cells (24) were labelled with pHrodo^TM^ Green STP ester (Thermo Fisher Scientific) as previously described and were used at the following respective concentrations which were determined to be non-saturating: 1 μM, 15 μg/mL and 50 000 cells/mL respectively. Bioparticles of pHrodo^TM^ Green-labelled *Escherichia coli* (*E. coli*) were purchased from Thermo Fisher Scientific and used at a concentration of 25 μg/mL. Cells were incubated with the labelled substrates for three hours unless otherwise indicated, and before being counterstained with Hoechst 33342 (5 μg/mL). Total green fluorescence intensity per cell was quantified on a CellInsight CX5 High Content Screening Platform. All conditions were assessed in triplicate. Unchallenged cells were used to measure background/autofluorescence. Internalization of fluorescein isothiocyanate (FITC) - labelled myelin was assessed similarly, except cells were washed with 4% trypan blue solution to quench extracellular fluorescence prior to imaging.

### Measurement of cytokine secretion

The following reagents were used to treat cells: lipopolysaccharide (LPS) from *E. coli* strain O127:B8 (100 ng/mL; Sigma Aldrich), Pam_3_CSK_4_ (100 ng/mL; Invivogen), R-FSL-(250 ng/mL; EMC Microcollection Gmbh), interferon gamma (IFNγ; 10 ng/mL; Peprotech) and adenosine triphosphate (ATP; 5 mM; Sigma Aldrich). Concentrations of interleukin-1beta (IL-1Π), interleukin-6 (IL-6), interleukin-10 (IL-10) and tumor necrosis factor (TNF) in cell supernatants were measured using the Human Inflammatory Cytokine Cytometric Bead Array Kit (BD Biosciences). Concentrations of chemokines in cell supernatants were measured using the LEGENDplex^TM^ Human Proinflammatory Chemokine Panel 1 (Biolegend). Readings were made on an Attune^TM^ Nxt Flow Cytometer.

### Western blotting

Cells were lysed on ice in a lysis buffer composed of 150 mM NaCl, 50 mM Tris-HCl pH 7.4, 1% Nonidet P-40, 0.1% sodium dodecyl sulfate (SDS) and 5 mM EDTA with protease and phosphatase inhibitors (Thermo Fisher Scientific). Cell lysates were centrifuged at 500 g for 30 minutes at 4°C to remove cellular debris. Proteins (25 μg/lane) were separated on SDS-polyacrylamide gels and transferred to polyvinylidene difluoride membranes (Bio-Rad Laboratories). Membranes were immunoblotted for CSF1R (1:250; #MAB3291, R&D Systems) and its phosphorylated form (Y723; 1:500; #3155, Cell Signaling), nuclear factor kappa B NF-κB (1:500; #8282, Cell Signaling) and its phosphorylated form (S536; 1:500; #3033, Cell Signaling), caspase-1 (1:500; #ab179515, Abcam), IL-1β (1:500; #12242, Cell Signaling), NLR family pyrin domain containing 3 (NLRP3; 1:500; #15101S, Cell Signaling) and GAPDH (1:5000; G8795, Sigma Aldrich) overnight at 4°C, and then with horse radish peroxidase-linked secondary antibodies (1:10000; Jackson Laboratory) for one hour. Bands were detected by enhanced chemiluminescence SignalFire^TM^ Plus ECL reagent (Cell Signaling) using a ChemiDoc Imaging System (Bio-Rad Laboratories). Image analysis was performed using ImageLab 6.0.1 software (Bio-Rad Laboratories).

### *CSF1R* sequencing

*CSF1R* pathogenic variant (c.2350G > A; p.V784M) in ALSP patient’s PBMCs and iPSCs was confirmed by Sanger sequencing. Following DNA extraction, touchdown polymerase chain reaction (PCR) was performed using the forward and reverse primers 5’ACGATACACATTCTCAGATCCTGG 3’ and 5’GTGTAGACACAGTCAAAGATGCTC 3’ respectively, for PBMCs, and 5’GGTAGGAGAAGGCCCAAGAC 3’ and 5’GGGATGACAGTCCCCAGTTA 3’, respectively, for iPSCs (designed using NCBI’s primer design tool, https://www.ncbi.nlm.nih.gov/tools/primer-blast/, and NM_005211.4 as reference sequence). The optimal annealing temperature for the primers was established at 54°C using the Tm Calculator provided by New England BioLabs (https://tmcalculator.neb.com/#!/main). The amplified deoxyribonucleic acid (DNA) sample corresponding to the PBMCs was sent to Genome Quebec, Montreal, Canada, for Sanger sequencing performed on an Applied Biosystems 3730xl DNA Analyzer (Thermo Fisher Scientific). For the iPSC-derived sample, sequencing was performed on an Applied Biosystems SeqStudio Genetic Analyzer (Thermo Fisher Scientific).

Sequencing of the entire region encoding the tyrosine kinase domain of CSF1R was performed for iMGL using ENST00000675795.1 as the reference transcript sequence. Following RNA extraction and reverse transcription, complementary DNA (cDNA) was subjected to amplification cycles using three sets of primers (Additional file 1: Figure S1A-B). The optimal annealing temperature for the primers was established at 52°C using the Tm Calculator provided by New England BioLabs (https://tmcalculator.neb.com/#!/main).

PCR products were purified using an ExoSAP-IT PCR product cleanup reagent (Thermo Fisher Scientific). DNA sequencing reactions were performed using a BigDye v3.1 cycle sequencing kit (Thermo Fisher Scientific) and followed by purification using a BigDye XTerminator Purification kit (Thermo Fisher Scientific). Sequencing was carried out on an Applied Biosystems SeqStudio Genetic Analyzer (Thermo Fisher Scientific).

### Magnetic resonance imaging (MRI)

Brain MRI was clinically performed in the 1.5T Philips MR scanner of the Montreal Neurological Institute, Montreal, Canada, with a protocol including 3D FLAIR T2-weighted images, axial T2-weighted images and diffusion weighted images.

### Migration assay

A Boyden chamber assay was carried out to assess cell migration toward adenosine diphosphate (ADP). Cells were plated on top compartments of Corning® Transwell® inserts with 8.0 μm pores (#3422), in nucleoside-free MEMα. Bottom compartments were filled with nucleoside-free MEMα containing vehicle or ADP (20 μM). When indicated, PSB0739 (20 μM) was added to both top and bottom compartments. Migration was quantified 1.5 hours later through Hoechst 33342 staining (5 μg/mL) of cells that crossed the inserts toward the lower compartment, using an EVOS M5000 Imaging System (Thermo Fisher Scientific).

### Assessment of lysosomal pH

Cells were incubated for one minute with 2.5 μM LysoSensor^TM^ Yellow/Blue DND-160 (Thermo Fisher Scientific). Fluorescence intensity was assessed using a SpectraMax® iD3 microplate reader, at excitation wavelengths of 329 and 384 nm, and emission wavelength of 540 nm. Unstained cells were used for background subtraction. Fluorescence intensity ratio (Ex 329:384) was calculated as an indicator of lysosomal pH as previously described (25). All conditions were assessed in triplicate. When indicated, cells were treated with 50 mM ammonium chloride (Sigma Aldrich) for one hour prior to the staining, throughout the staining, and during fluorescence intensity measurement.

### Assessment of lysosomal protease activity

Cells were incubated with DQ^TM^ red bovine serum albumin (BSA) following the manufacturer’s recommendation for 24 hours. Total red fluorescence intensity per cell was quantified using a CellInsight CX5 High Content Screening Platform. All conditions were assessed in triplicate. Unchallenged cells were used to measure background/autofluorescence. When indicated, cells were concomitantly treated with 100 nM Bafilomycin A1 (Thermo Fisher Scientific).

### Statistical analyses

Statistical analyses were performed using GraphPad Prism 9.0 software. A t-test was used to compare the mean of two groups of data. A one-way analysis of variance (ANOVA) was used to compare the mean of three or more groups of data. When the assumptions of a t-test or a one-way ANOVA were not met, a Mann-Whitney test or a Kruskal-Wallis test was used instead, respectively. P-values (‘p’) were adjusted using appropriate post hoc tests following one-way ANOVA/Kruskal-Wallis tests. A two-way ANOVA followed by Sidak multiple comparison tests was used to compare two groups of data with multiple variables. Mean and standard error of the mean (SEM) of biological replicates (‘n’) are plotted in all graphs unless otherwise indicated. A p < 0.05 was considered statistically significant. Primary microglia obtained from independent donors and iMGL generated at different points in time were considered biological replicates.

## Results

### The 2.9 protocol yields higher number of pure, adherant iMGL compared to the 2.0 protocol

The newly devised 2.9 protocol consisted of two steps (Figure 1A): 1) generation of iHPCs from iPSCs identically to McQuade *et al*.’s 2.0 protocol using a commercially available kit, and 2) differentiation of iHPCs into iMGL using a modified medium formulation from the 2.0 protocol (Table 1). MEMα containing physiological concentration of glucose was used as the base of the 2.9 microglia differentiation medium, instead of DMEM/F12 in the 2.0 medium. The growth factor granulocyte-macrophage colony-stimulating factor (GM-CSF) at low concentration has been previously shown to be beneficial in increasing iMGL cell yield (26) and was therefore incorporated as a mitogenic factor, in addition to IL-34, M-CSF and TGF-β1. While C-X3-C motif chemokine ligand 1 (CX3CL1) and cluster of differentiation 200 (CD200) had been used in the 2.0 protocol as modulators of microglia function, CD200 was omitted in the 2.9 protocol as the expression of its receptor is almost absent in human microglia (15, 27). Finally, monothioglycerol and N2 supplements were omitted, as they have been shown to provide no benefit in microglia identity acquisition (26).

**Figure 1.**
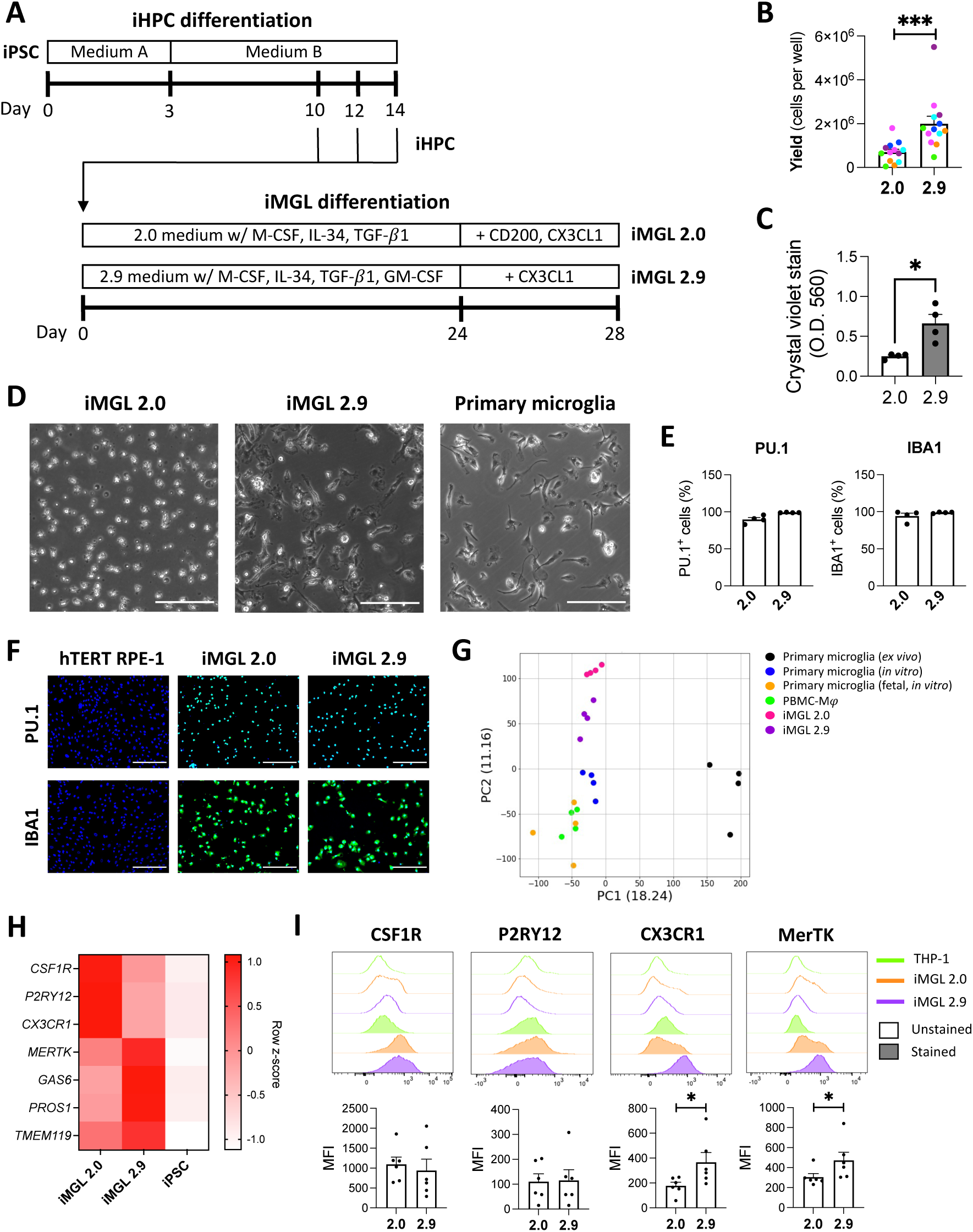
Characterization of iMGL 2.9. (A) Schematic of the 2.0 and 2.9 protocols. (B-H) iMGL 2.0 and 2.9 differentiation were carried out side-by-side from the same iPSC lines. (B) Viable iMGL cell yield assessed by trypan blue exclusion assay. Cell numbers per well of a 6-well plate are presented. A t-test was performed. n = 13 differentiation batches from 6 iPSC lines. *** p < 0.001. (C) Crystal violet assay. A Mann-Whitney test was performed. n = 4 differentiation batches from 4 iPSC lines, * p < 0.05, O.D. = optical density. (D) Phase contrast images of iMGL 2.0, iMGL 2.9 and primary human microglia. Scale bar = 150 𝜇𝑚. (D) Quantification of PU.1- and IBA1-positivity by immunostaining. n = 4 differentiation batches from 4 iPSC lines. (E) Representative images of PU.1 (green) and IBA1 (green) immunostaining of iMGL 2.0 and 2.9 counterstained with Hoescht 33342 (blue). Immunostaining of hTERT RPE-1 cells are shown as negative controls. Scale bar = 200 𝜇𝑚. (F) PCA plot of RNA-sequencing data. (G) Heatmap showing key microglia marker expression assessed by qRT-PCR in iMGL 2.0, iMGL 2.9 and iPSCs. n = 6 differentiation batches from 6 iPSC lines. (H) Flow cytometry assessment of cell surface microglia marker expression in iMGL 2.0 and 2.9. Data from THP-1 cells are shown as negative controls. Mann-Whitney tests were performed. n = 6 differentiation batches from 6 iPSC lines, * p < 0.05. MFI = median fluorescence intensity.

From the same starting number of iHPCs, the new 2.9 protocol resulted on average in ∼3 times higher number of viable iMGL compared to the 2.0 protocol (Figure 1B), with some variability in yield improvement observed between different iPSC lines (between 2 to 8-fold increase; Additional file 1: Figure S2A). In contrast to the 2.0 protocol that yielded loosely adherent and floating cells, the 2.9 protocol resulted in more tightly adherent cells of elongated, amoeboid or ramified morphologies akin to primary cells (Figure 1C-D, Additional file 1: Figure S2B). Occasional formation of multinucleated giant cells could be observed (Additional file 1: Figure S3A), but those could be eliminated through the selective harvest of mononuclear cells using a 2 mM EDTA solution (Additional file 1: Figure S3B). Immunostaining revealed iMGL 2.9 to be ∼99% and ∼98% positive for the respective myeloid markers PU.1 and IBA1 (Figure 1E-F; the neuroepithelial cell line hTERT RPE-1 was used as a negative control).

RNA-seq revealed the transcriptomes of iMGL 2.0 and 2.9 to greatly differ from that of iPSCs from which they were generated, and closely resembled *in vitro* cultured primary microglia and macrophages (Additional file 1: Figure S4). Transcriptomes of iMGL generated using either the 2.0 or 2.9 protocols slightly differed from cultured (*in vitro*) and freshly isolated (*ex vivo*) primary microglia, but iMGL 2.9 showed a higher transcriptomic similarity to *in vitro* and *ex vivo* primary microglia compared to iMGL 2.0 (Figure 1G). DEG analysis revealed genes more highly expressed in iMGL 2.0 over 2.9 to be enriched in processes related to cell division (Additional file 1: Figure S5A). Genes more highly expressed in iMGL 2.9 over 2.0 were involved in microglial cell activation (*e.g.* triggering receptor expressed on myeloid cell 2 or *TREM2*, chemokine ligand 3 or *CCL3*, metalloproteinase 8 or *MMP8*…), chemotactic attraction of leukocytes and complement system (Additional file 1: Figure S5A). A number of macrophage-specific markers such as myeloperoxidase (*MPO*) were confirmed to be lowly expressed in both iMGL 2.0 and 2.9, compared to PBMC-Mχπ (Additional file 1: Figure S5B). Cluster of differentiation 36 (*CD36*) was previously described as a marker of fetal microglia (28) and was more highly expressed on cultured fetal microglia and PBMC-Mχπ than postnatal microglia or iMGL 2.0 and 2.9 (Additional file 1: Figure. S5B). Genes that are known to be more highly expressed in microglia over macrophages such as *P2RY12*, G-protein coupled receptor 34 (*GPR34*) or sialic acid binding Ig-like lectin 10 (*SIGLEC10*) were more highly expressed in iMGL 2.9 compared to PBMC-Mχπ (Additional file 1: Figure S5B). Assessment of homeostatic microglia markers revealed some markers such as *CSF1R*, *P2RY12* and *CX3CR1* to be lower in iMGL 2.9 compared to iMGL 2.0, whereas other markers such as *MERTK*, growth arrest-specific 6 (*GAS6*) and protein S (*PROS1*) were higher in iMGL 2.9 (Figure 1H). However, flow cytometry assessment of cell surface expression revealed CX3CR1 and MerTK to be higher in iMGL 2.9 compared to iMGL 2.0, and CSF1R and P2RY12 to be similar between the two protocols (Figure 1I; the monocytic cell line THP-1 was used as a negative control). Overall, our findings imply that the 2.9 protocol results in an improved yield of more adherent microglia-like cells with higher transcriptomic similarity to primary microglia compared to the original 2.0 protocol.

### iMGL 2.9 are better phagocytes than iMGL 2.0

*MERTK*, *GAS6* and *PROS1*, which were found to be more highly expressed in iMGL 2.9 compared to iMGL 2.0, are all well known for their involvement in phagocytic processes (15, 23, 29). In addition, DEG analysis revealed a number of other genes implicated in phagocytosis, such as genes encoding Fcγ receptors (Figure 2A), to be more highly expressed in iMGL 2.9 compared to iMGL 2.0. Consistently, phagocytosis assay revealed a higher uptake of pHrodo^TM^ Green-labelled myelin and α-synuclein fibrils (both mediated by MerTK (23, 30)), and IgG-opsonized red blood cells (mediated by Fcγ receptors) by iMGL 2.9 compared to iMGL 2.0 (Figure 2B-C). No difference in the extent of *E. coli* uptake was observed between iMGL 2.0 and 2.9 (Figure 2B-C). All substrates were internalized by iMGL 2.9 at a comparable level to primary microglia (Figure 2B-C), but this comparison could not be systematically carried out due to the limited availability of primary microglia. Overall, iMGL generated using the 2.9 protocol are more efficiently able to phagocytose a variety of substrates better than cells generated using the original 2.0 protocol.

**Figure 2.**
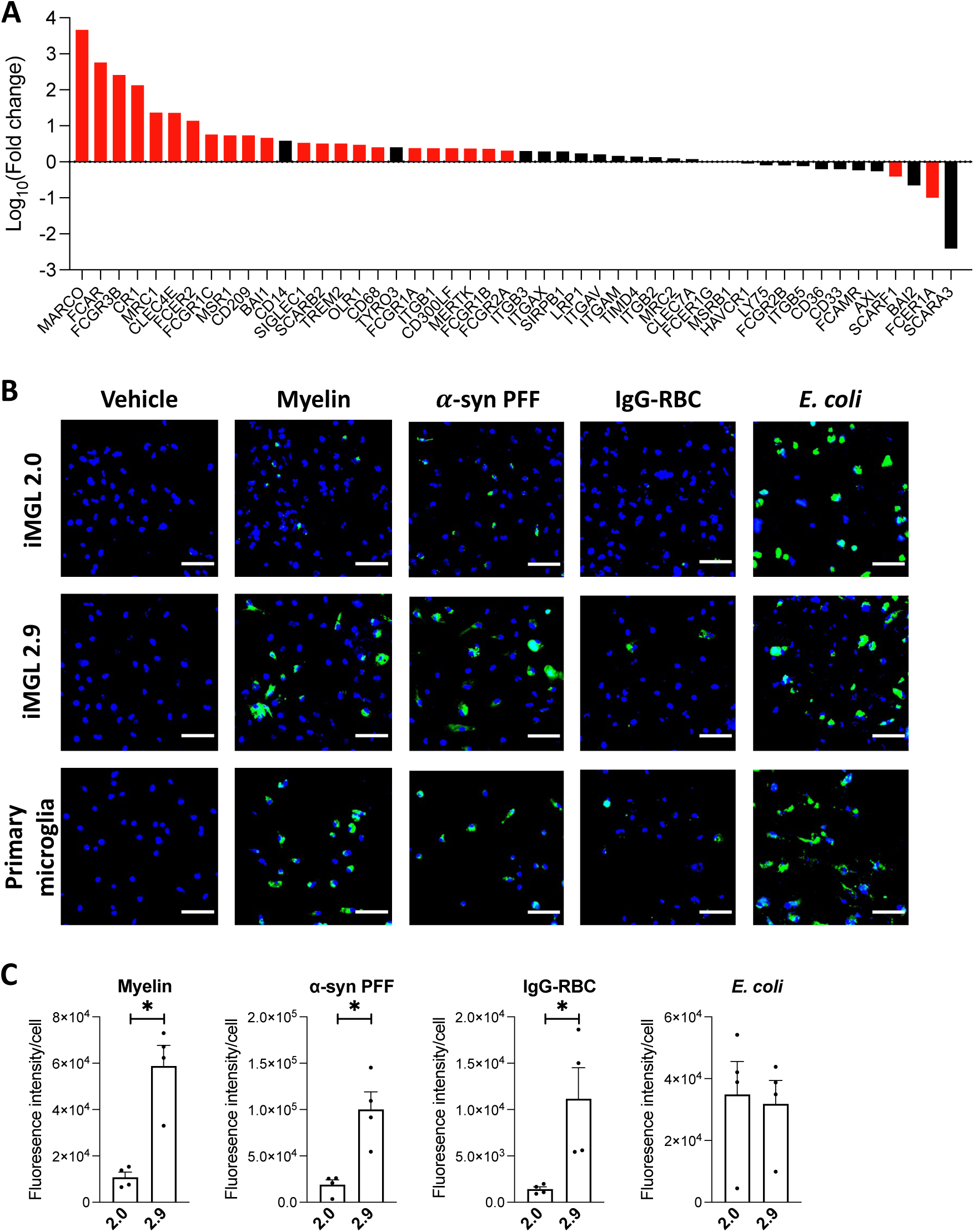
Phagocytic competence of iMGL 2.9. iMGL 2.0 and 2.9 were differentiated side-by-side from the same iPSC lines. (A) Expression of genes encoding phagocytosis/scavenger receptors or positive regulators of phagocytosis in iMGL 2.9 compared to 2.0. Red bars represent genes that were significantly different (adjusted p < 0.05) between iMGL 2.0 and 2.9 whereas black bars represent genes that were not significantly different (n = 4 differentiation batches from 4 iPSC lines). (B-C) iMGL 2.0, iMGL 2.9 and primary human microglia were treated side-by-side with vehicle or pHrodo^TM^ Green-labelled myelin, α-synuclein preformed fibrils (α-syn PFFs), opsonized red blood cells (IgG-RBC) or *E. coli* for three hours and then counterstained with Hoescht 33342. (B) Representative fluorescence images. Scale bar = 50 𝜇𝑚. (C) Quantification of green fluorescence intensity per cell. Mann-Whitney tests were performed. n = 4 differentiation batches from 4 iPSC lines, * p < 0.05.

### LPS elicits a functional TLR4 response from iMGL 2.9 but not iMGL 2.0

A major issue previously noted with iMGL differentiated using the 2.0 protocol is their poor inflammatory response to LPS (15, 31), a widely used inflammatory stimulus that agonizes TLR4. Measurement of cytokine secretion following LPS treatment revealed iMGL 2.9 and primary microglia, but not iMGL 2.0, to secrete significantly higher amounts of IL-6, TNF and IL-10 (Figure 3A). This was associated with increased phosphorylation of the transcription factor NF-κB regulating cytokine expression following LPS treatment in iMGL 2.9 (Figure 3B-C). Accordingly, flow cytometry (Figure 3D) and qRT-PCR (Additional file 1: Figure. S6) assessment of TLR4 and its co-receptor CD14, essential for LPS recognition (32), revealed CD14 expression to be significantly higher in iMGL 2.9 compared iMGL 2.0. Expression of lymphocyte antigen 96 (*LY96*), encoding the co-receptor of TLR4 MD2 also essential for LPS recognition (32), was also more highly expressed in iMGL 2.9 compared to iMGL 2.0 as quantified by qRT-PCR (Additional file 1: Figure S6). MD2 cell surface expression was not investigated due to the unavailability of flow cytometry-validated antibodies. RNAseq analysis revealed no difference in the expression of pathogen recognition receptors and their co-receptors between iMGL 2.0 and 2.9, with the exception of *CD14* (Figure 3E). Interestingly, substitution of DMEM/F12 by MEMα in the 2.0 microglia differentiation medium was sufficient to increase the inflammatory response of iMGL to LPS, but this was further promoted by the addition of GM-CSF (Additional file 1: Figure S7). iMGL 2.9 also showed functional inflammatory response to agonists of other TLRs such as Pam_3_CSK_4_ or R-FSL-1. (Additional file 1: Figure S8A). Secretion of a wide array of chemokines by iMGL 2.9 was observed following LPS or IFNγ treatment, albeit with some discrepancy in secretory pattern compared to primary microglia (Additional file 1: Figure S8B).

**Figure 3.**
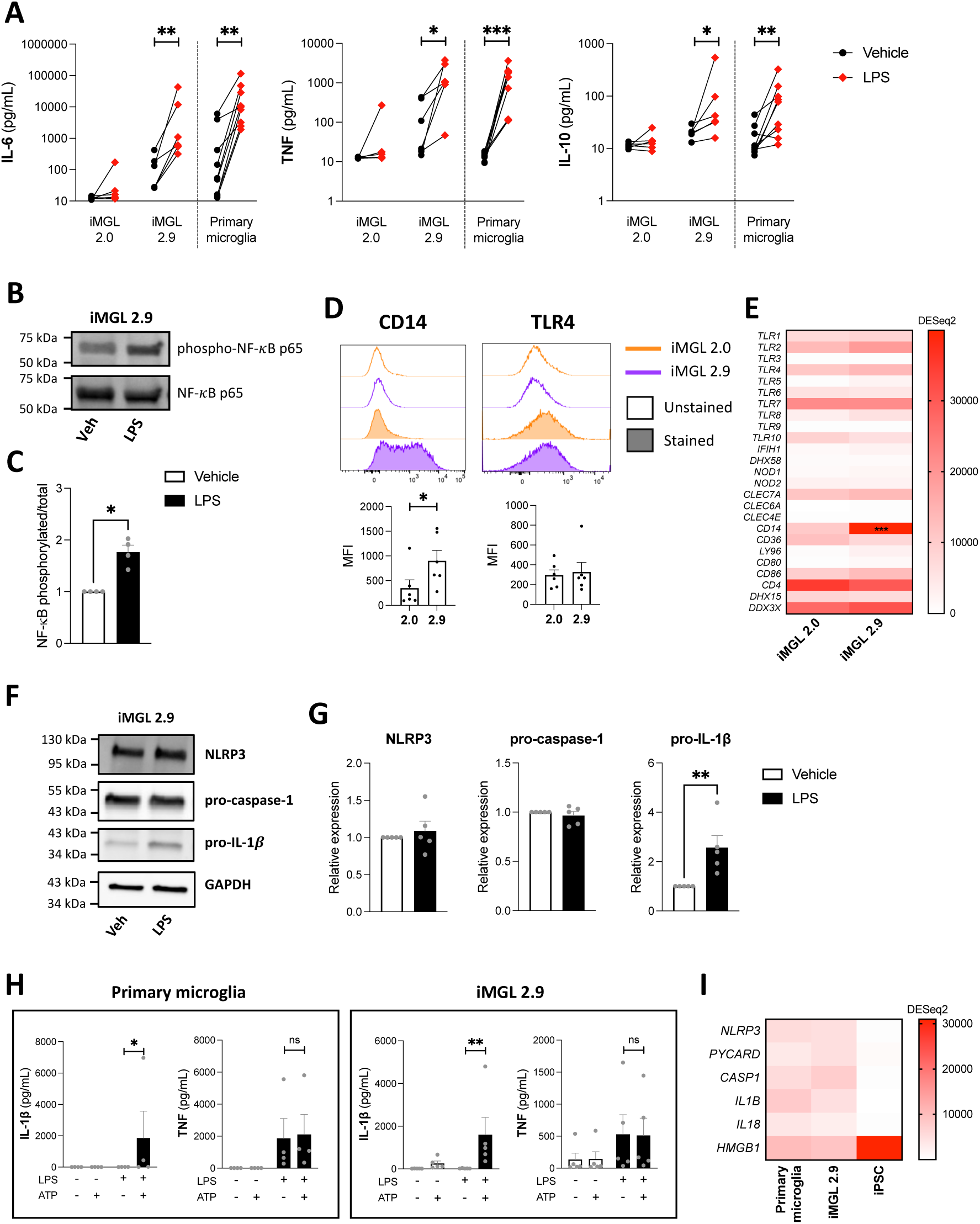
TLR4 signaling response of iMGL 2.9 following LPS treatment. iMGL 2.0 and 2.9 were differentiated side-by-side from the same iPSC lines. (A) IL-6, TNF and IL-10 concentrations in supernatants from iMGL 2.0, iMGL 2.9 and primary human microglia treated with vehicle or LPS (100 ng/mL) for 24 hours. Mann-Whitney tests were performed. n = 5 differentiation batches from 5 iPSC lines for iMGL, n = 9 donors of primary microglia, * p < 0.05, ** p < 0.01, *** p < 0.001. (B) Representative Western blotting and (C) quantification of NF-κB in iMGL 2.9 treated with vehicle or LPS (100 ng/mL) for 1.5 hours. A Mann-Whitney test was performed. n = 4 differentiation batches from 4 iPSC lines, * p < 0.05. (D) Flow cytometry assessment of cell surface CD14 and TLR4 expression in iMGL 2.0 and 2.9. Mann-Whitney tests were performed. n = 6 differentiation batches from 6 iPSC lines, * p < 0.05. (E) Heatmap showing expression of genes encoding pattern recognition receptor and their co-receptors. A two-way ANOVA was performed. n = 4 differentiation batches from 4 iPSC lines, *** p < 0.001 vs iMGL 2.0. (F) Representative Western blotting and (G) quantification of inflammasome components in iMGL 2.9 treated with vehicle or LPS (100 ng/mL) for 24 hours. Mann-Whitney tests were performed. n = 5 differentiation batches from 5 iPSC lines, * p < 0.05. (H) IL-1ý and TNF concentrations in cell supernatants of human primary microglia and iMGL 2.9 treated with vehicle or LPS (100 ng/mL) for 24 hours, followed by ATP (5 mM) or not for 30 minutes. Kruskal-Wallis tests followed by Dunn’s multiple comparison tests were performed. n = 4 donors of primary microglia and n = 5 differentiation batches from 5 iPSC lines, * p < 0.05, ** p < 0.01, ns = non-significant. (I) Heatmap showing the expression of genes encoding NLRP3 inflammasome components and their substrates. n = 4 primary microglia donors, 4 differentiation batches from 4 iPSC lines for iMGL, and 4 iPSC lines.

The inflammasome is a stimulus-induced multiprotein complex of the innate immune system that have been linked to a variety of diseases, including neurodegenerative diseases (33–36). iMGL 2.9 had a functional inflammasome system, as evidenced by increased IL-1ý, but not TNF release upon ATP treatment, when cells were primed with LPS to increase IL-1β protein expression (Figure 3F-H). Similarly, cultured primary microglia also showed enhanced IL-1β secretion following LPS priming prior to ATP induction of the inflammasome (Figure 3H). iMGL 2.9 and primary microglia in culture had similar basal expression of genes encoding NLRP3 inflammasome components and their substrates, and distinctively differed from iPSCs used to derive the iMGL (Figure 3I). Overall, our findings imply that iMGL derived using the 2.9 protocol are better suited for studying microglial inflammatory activities compared to those derived using the 2.0 protocol.

### The c.2350G > A *CSF1R* variant causes ALSP

The primary objective of the current study was to develop an *in vitro* tool to study *CSF1R* variants associated with ALSP in human microglia. A 54-year-old male patient of Croatian origin (Figure 4A subject II.3) was diagnosed with ALSP in 2020. A missense mutation (c.2350G > A) resulting in the substitution of a highly conserved valine residue by a methionine residue in the tyrosine kinase domain of CSF1R (p.V784M) had been previously described in his deceased sister (Figure 4A subject II.2) who had also been clinically diagnosed with ALSP (37). His mother (Figure 4A subject I.1) is also a carrier of the same *CSF1R* variant and presented with a well-controlled bipolar disease, but no neurological symptoms or dementia. His family history was also positive for schizophrenia in the oldest sister (Figure 4A subject II.1), who refused genetic testing. A few months prior to the diagnosis, the patient manifested mood swings, personality changes, and trouble performing common tasks at work. Upon assessment at the Montreal Neurological Institute-Hospital, his neurological examination revealed anxiety, mild dysmetria and gait apraxia. Brain MRI documented extensive bilateral white matter abnormalities predominant in the frontal and parietal lobes, more prominent on the right hemispheres (Figure 4B-D). The corpus callosum was also affected at the level of the genu and anterior portion of the body (Figure 4B). On diffusion-weighted images, few foci of diffusion restriction in the corona radiata and centrum semiovale were detected bilaterally (Figure 4E). Genetic sequencing of *CSF1R* documented the presence of the heterozygous c.2350G > A (p.V784M) pathogenic variant in the patient, confirming the diagnosis of ALSP (Figure 4F). Given the clear association of this c.2350G > A variant with clinical diagnosis of ALSP, the patient’s PBMCs were collected in order to generate iMGL. The patient requested and received medical assistance in dying at the age of 57.

**Figure 4:**
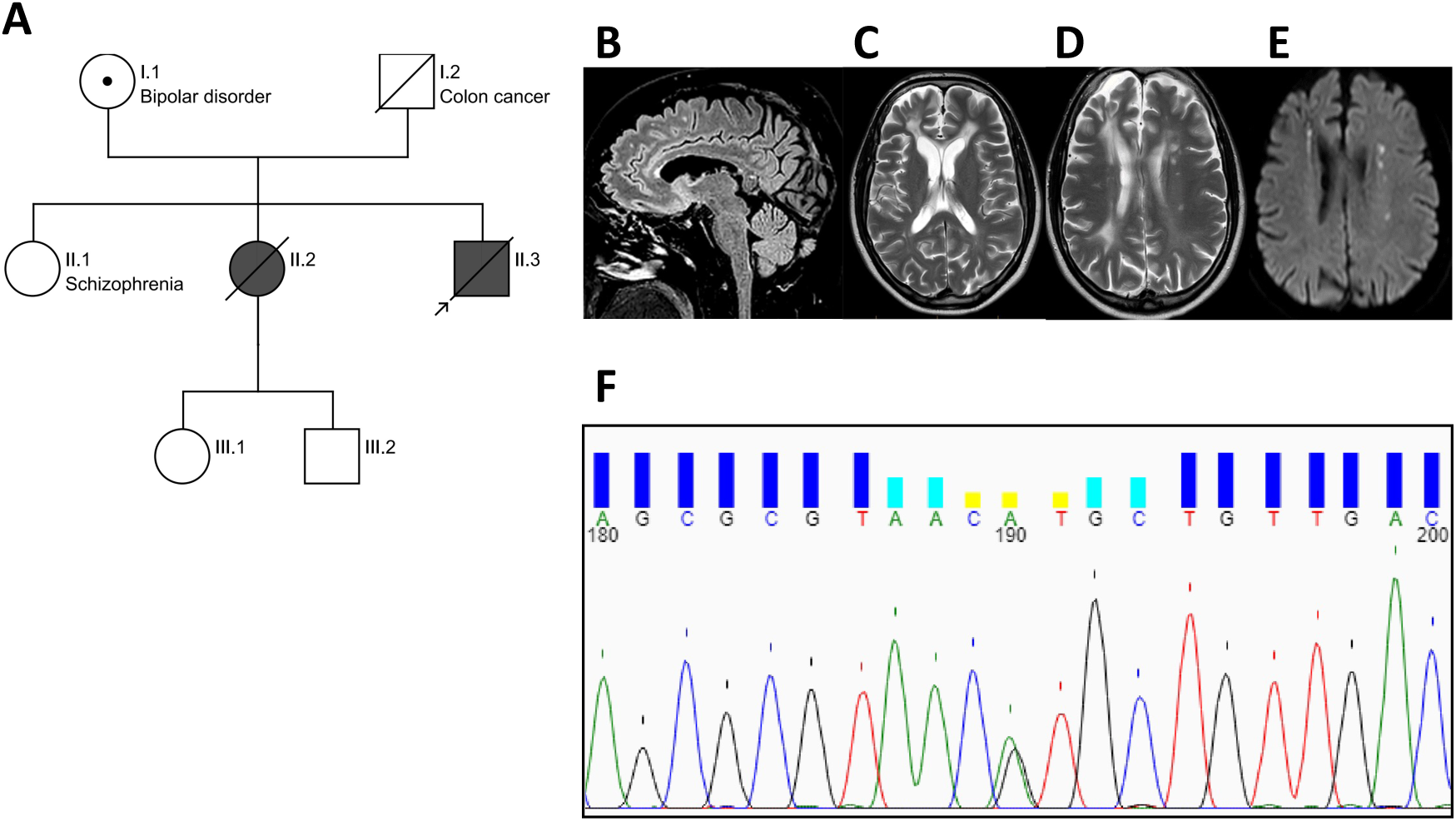
Pedigree of the proband’s family, MRI images, and genotyping data. (A) Pedigree of the proband’s family (prepared via https://cegat.com/). Shading indicates carriers of the c.2350G >A *CSF1R* variant with ALSP diagnosis. Symbol with a dot indicates a carrier without clinical manifestations of ALSP. (B) Sagittal FLAIR T2-weighted MR image showing thinning and hyperintense signal of the anterior portion of the body of the corpus callosum (red arrow). (C-D) Axial T2-weighted MR images showing the presence of bilateral multifocal and confluent lesions in the frontal lobes and multifocal lesions in the left fontal and parietal lobe with areas of restricted diffusion in the diffusion weighted image (E). (F) Chromatogram of PBMC-derived DNA showing heterozygous c.2350G > A *CSF1R* variant (shown here as position 190).

### The 2.9, but not the 2.0 protocol, allows the generation of viable iMGL from an ALSP patient harboring a heterozygous *CSF1R* pathogenic variant

Patient’s PBMCs were first reprogrammed into iPSCs that expressed the pluripotency markers Nanog, Tra-1-60, SSEA-4 and OCT3/4 (Additional file 1: Figure S9A). Presence of a c.2350G > A variant in the *CSF1R* gene was confirmed by Sanger sequencing (Additional file 1: Figure S9B). Karyotyping and qPCR-based screening did not reveal any genomic abnormality (Additional file 1: Figure S9C-D). The patient line will be referred to as “ALSP-CSF1R”. Differentiation of ALSP-CSF1R iPSCs into iHPCs resulted in round floating CD43+ cells (Additional file 1: Figure S10A-B). A fraction of these CD43+ cells also expressed the hematopoietic stem cell marker CD34 (Additional file 1: Figure S10B) as expected (4). While the 2.0 protocol failed to generate any viable microglial cells from the ALSP-CSF1R iHPCs, the 2.9 protocol successfully induced their microglial differentiation (Figure 5A-B). Interestingly, addition of GM-CSF to the 2.0 microglia differentiation medium and the substitution of DMEM/F12 with MEMα, but neither of these medium modifications alone, resulted in the robust generation of viable iMGL from ALSP-CSF1R iHPCs (Additional file 1: Figure S11). When the 2.9 protocol was used, the final cell yield was significantly lower with the ALSP-CSF1R line compared to healthy control lines differentiated side-by-side (Figure 5C). ∼100% and ∼99% of the ALSP-CSF1R iMGL expressed PU.1 and IBA1, respectively (Figure 5D), indicative of successful myeloid differentiation. The cells showed a time-dependent increase in the expression of microglia marker genes such as *CX3CR1*, *GAS6* and transmembrane protein 119 (*TMEM119*) throughout their differentiation, however *P2RY12* expression was observed to decline over the latter half of the differentiation period (Figure 5E). This resulted in significantly lower expression of *P2RY12* in ALSP-CSF1R iMGL compared to healthy controls (Figure 5E).

**Figure 5.**
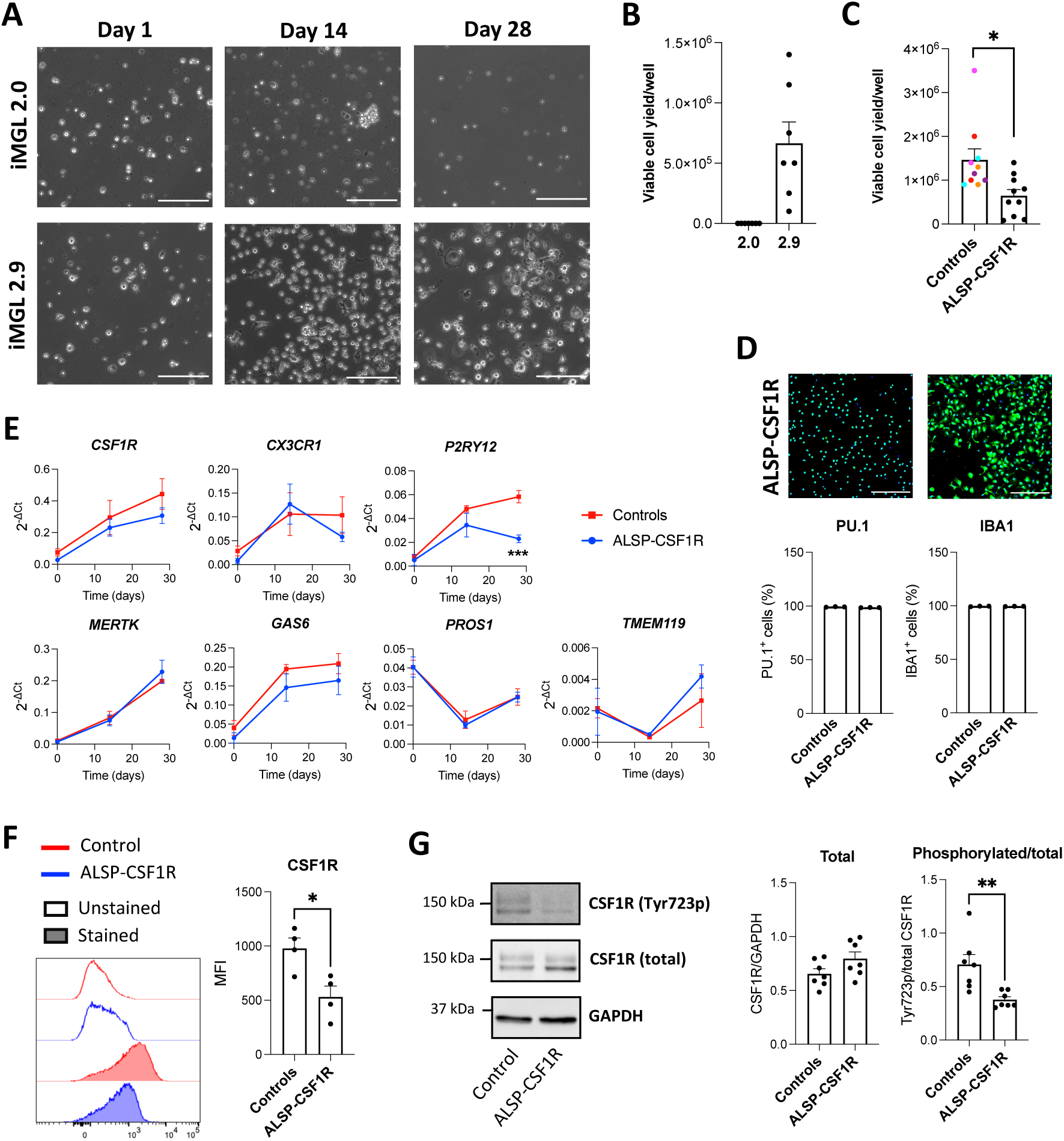
Derivation of microglia-like cells from the ALSP-CSF1R patient using the 2.9 protocol. (A) Phase contrast images of ALSP-CSF1R iPSC line differentiated into iMGL using the 2.0 or 2.9 protocol. Scale bar = 150 𝜇𝑚. (B) Viable cell yield per well of a 6-well plate assessed by trypan blue exclusion assay following 2.0 and 2.9 differentiation of the ALSP-CSF1R iPSCs into iMGL. n = 7 differentiation batches. (C) Viable cell yield per well of a 6-well plate assessed by trypan blue exclusion assay following side-by-side differentiation of healthy control lines (n = 4 differentiation batches from 4 iPSC lines) and the ALSP-CSF1R line (n = 4 differentiation batches) using the 2.9 protocol. A t-test was performed, * p < 0.05. (D) Quantification of PU.1-and IBA1-positive cells by immunostaining. n = 4 batches from 4 healthy control iPSC lines and 4 batches of a single ALSP-CSF1R line, differentiated side-by-side using the 2.9 protocol. (E) qRT-PCR assessment of microglia marker expression on day 0, 14 and 28 of microglial differentiation. n = 4 batches from 4 healthy control iPSC lines and 4 batches of a single ALSP-CSF1R line, differentiated side-by-side using the 2.9 protocol. Two-way ANOVA were performed, followed by Sidak’s post hoc tests. *** p < 0.001. (F) Flow cytometry assessment of CSF1R cell surface expression. A t-test was performed. n = 4 batches from 4 healthy control iPSC lines and 4 batches of a single ALSP-CSF1R line, differentiated side-by-side using the 2.9 protocol. * p < 0.05. (G) Western blot assessment of CSF1R and its tyrosine 723-phosphorylated form, and GAPDH. A t-test was performed. n = 7 batches from 4 healthy control iPSC lines and 7 batches of a single ALSP-CSF1R line, differentiated side-by-side using the 2.9 protocol. ** p < 0.01.

ALSP-CSF1R iMGL expressed the c.2350G > A *CSF1R* variant, whereas none of the iMGL used as controls had pathogenic variants in the region encoding CSF1R tyrosine kinase domain (Additional file 3: Table S2). *CSF1R* mRNA (Figure 5E) and protein (Figure 5G) expression in ALSP-CSF1R iMGL was no different from healthy controls, yet cell surface expression of CSF1R measured by flow cytometry was significantly lower (Figure 5F), indicative of a defective cell surface trafficking or recycling of CSF1R in the patient-derived cells. This was consistent with a previous report that ALSP-associated mutant CSF1R accumulate in Golgi-like perinuclear regions, resulting in reduced cell surface expression (38). Assessment of CSF1R phosphorylation at tyrosine 723 revealed activation of CSF1R in ALSP-CSF1R iMGL to be lower compared to controls (Figure 5G), in line with previous observations that ALSP-associated variants in *CSF1R* impairs M-CSF-induced autophosphorylation (39, 40). Overall, yield assessment and molecular characterization revealed that ALSP-CSF1R iPSCs could be successfully differentiated into iMGL through the 2.9 protocol, but not the original 2.0 protocol.

### ALSP-CSF1R iMGL presents phenotypic alterations compared to healthy control iMGL

*P2RY12* encodes a purinergic receptor important for chemotactic migration of microglia toward nucleotides released by dead cells (41). Since *P2RY12* expression was observed to be lower in ALSP-CSF1R iMGL compared to controls, migratory activity was assessed using a Boyden chamber assay. While healthy control iMGL showed significant migration toward ADP (∼7-fold increase in migrating cells, p = 0.0103) that was blocked by the P2RY12 antagonist PSB0739, ALSP-CSF1R iMGL showed minimal migration toward ADP (∼2-fold increase in migrating cells, p = 0.0876; Figure 6A-B).

**Figure 6.**
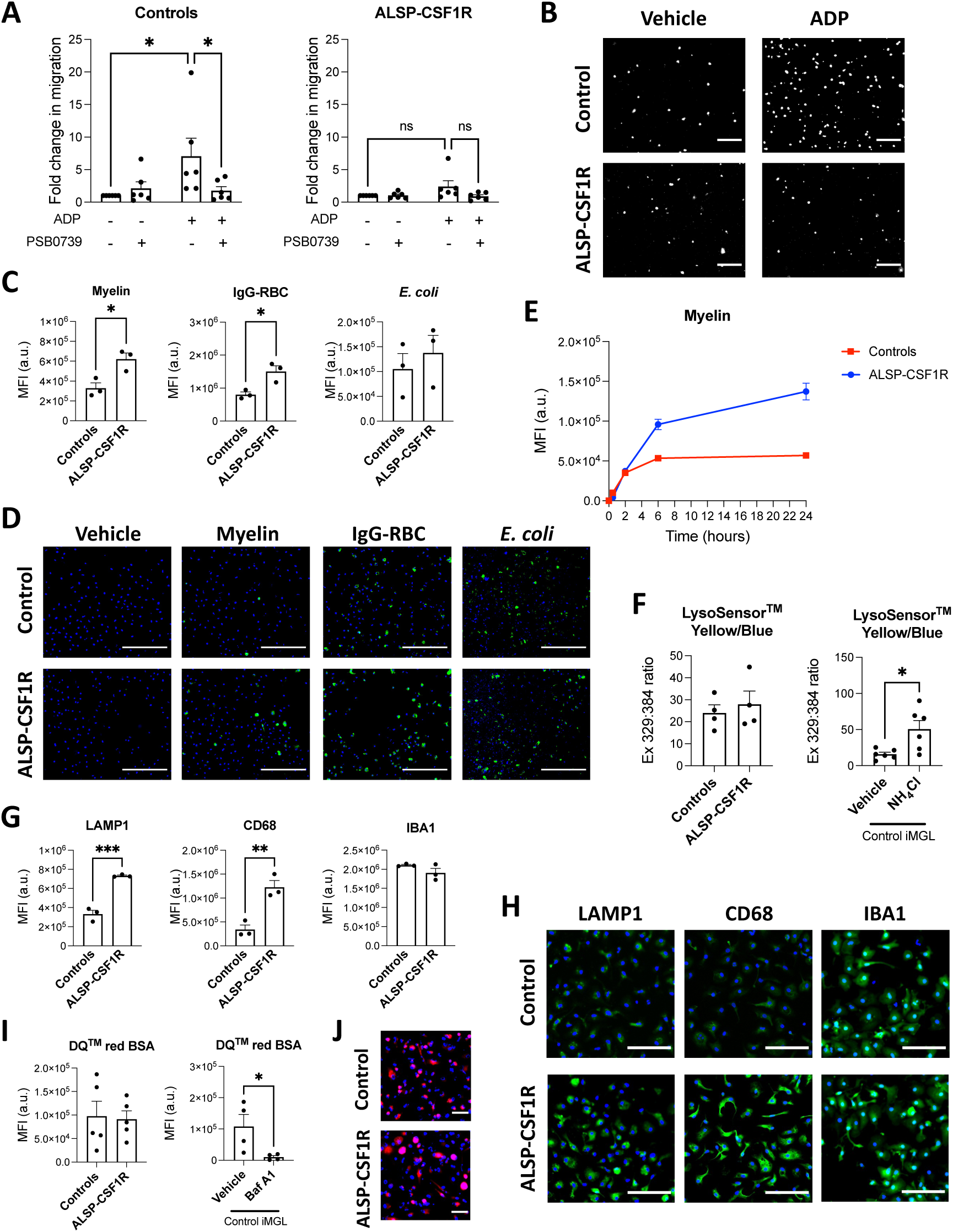
Functional phenotype of iMGL derived from the ALSP-CSF1R patient. All iMGL were generated using the 2.9 protocol. Quantification (A) and images (B) of iMGL migratory activity toward ADP assessed by Boyden chamber assay. Cells were concomitantly treated or not with PSB0739. Kruskal-Wallis tests followed by Dunn’s multiple comparison tests were performed. n = 6 batches from 4 healthy control iPSC lines and 6 batches of a single ALSP patient line, differentiated side-by-side. White = Hoechst 33342, scale bar = 75 μm in (B). Quantification of green fluorescence intensity per cell (C) and representative images (D) of iMGL exposed to vehicle or pHrodo^TM^ Green-labelled myelin, opsonized red blood cells (IgG-RBC) or *E. coli* for three hours and then counterstained with Hoescht 33342. T-tests were performed. n = 3 batches from 3 healthy control iPSC lines and 3 batches of a single ALSP patient line, differentiated side-by-side, * p < 0.05. Scale bar = 250 𝜇𝑚 in (D). (E) Timecourse of pHrodo^TM^ Green-labelled myelin uptake by control and ALSP-CSF1R iMGL. Mean +/-standard deviation of three technical replicates are presented. MFI = median fluorescence intensity, a.u. = arbitrary unit. (F) Assessment of lysosomal pH using LysoSensor^TM^ Yellow/Blue. Cell treatment with 50 mM ammonium chloride (NH_4_Cl) was used as a positive control. Left panel: n = 4 batches from 4 healthy control iPSC lines and 4 batches of a single ALSP patient line, differentiated side-by-side. Right panel: n = 6 batches from 4 healthy control iPSC lines. T-tests were performed, * p < 0.05. Quantification of mean fluorescence intensities (MFI; G) and representative images (H) of LAMP1, CD68 and IBA1 immunostaining of iMGL. T-tests were performed. n = 3 batches from 3 healthy control iPSC lines and 3 batches of a single ALSP patient line, differentiated side-by-side, ** p < 0.01, *** p < 0.001. Scale bar = 100 𝜇𝑚 in (H). (I-J) Assessment of lysosomal protease activity using DQ^TM^ red BSA. (I) Cell treatment with bafilomycin A1 (Baf A1, 100 nM) was used as a control. MFI = mean fluorescence intensity, a.u. = arbitrary unit. Left panel: n = 5 batches from 4 healthy control iPSC lines and 5 batches of a single ALSP patient line, differentiated side-by-side. Right panel: n = 4 batches from 4 healthy control iPSC lines. T-tests were performed, * p < 0.05. (J) Representative fluorescence images. Scale bar = 50 𝜇𝑚.

Migration of microglia toward ADP is important for the phagocytic clearance of dead cells and debris. Surprisingly, phagocytosis assays using pHrodo^TM^ Green-labelled substrates revealed a higher uptake of myelin and IgG-opsonized red blood cells by ALSP-CSF1R iMGL compared to healthy controls (Figure 6C-D). Slightly higher uptake of *E. coli* was also observed in patient-derived cells, although this did not reach statistical significance (Figure 6C-D). Time course experiments using pHrodo^TM^ Green-labelled myelin pointed toward enhanced phagocytic capacity, rather than enhanced phagocytic rate, of the ALSP-CSF1R iMGL compared to controls (Figure 6E). pHrodo^TM^ Green is a pH-sensitive dye that allows the selective quantification of labelled targets that reaches acidic compartments of the cells upon phagocytosis (42, 43). Uptake assay using FITC-labelled myelin resulted in similar observations as with pHrodo^TM^ Green-labelled myelin (Additional file 1: Figure S12), excluding lysosomal pH as a factor influencing phagocytosis assay results. In line with this, lysosomal pH assessment using the ratiometric LysoSensor^TM^ Yellow/Blue probe revealed no difference in lysosomal pH between control and ALSP-CSF1R iMGL (Figure 6F). Treatment of iMGL with ammonium chloride resulted in increased lysosomal pH as expected (Figure 6F). Immunostaining intensities of the lysosomal markers LAMP1 and CD68 were higher in ALSP-CSF1R iMGL compared to healthy controls (Figure 6G-H). In contrast, staining intensities of the calcium-binding protein IBA1 were found to be similar between ALSP-CSF1R and control cells (Figure 6G-H). Overall, these data suggest that lysosomal content is higher in ALSP-CSF1R iMGL compared to controls, which could account for a higher phagocytic capacity of the patient-derived cells. DQ^TM^ red BSA is a labelled BSA that emits fluorescence upon BSA proteolysis in lysosomes. No difference in DQ^TM^ red BSA proteolysis was observed between control and ALSP-CSF1R iMGL over the course of 24 hours (Figure 6I). The lysosomal acidification inhibitor bafilomycin A1 significantly reduced DQ^TM^ red BSA proteolysis (Figure 6I). This indicates that despite higher internalization and lysosomal storage of phagocytosed materials, lysosomal protease activities are not higher in ALSP-CSF1R iMGL compared to controls. In sum, these findings suggest that the heterozygous c.2350G > A *CSF1R* variant is associated with phenotypic alterations of microglia including reduced chemotaxis toward ADP and higher phagocytic capacity owing to a larger lysosomal storage capacity.

## Discussion

To summarize, we have developed a new protocol for microglia derivation from iPSCs, resulting in a significant improvement in cell yield, adhesive properties, protein expression of select microglia markers, phagocytic activities and inflammatory response compared to the original protocol published by McQuade *et al.* in 2018. Transcriptomic profile of the resulting iMGL also more closely resembled that of *ex vivo* and *in vitro* primary human microglia. This new protocol was applied to generate iMGL from a *CSF1R*-mutated ALSP patient. Characterization of the ALSP-CSF1R iMGL carried out in the current study revealed lower CSF1R cell surface expression and autophosphorylation compared to control iMGL, as well as phenotypic alterations affecting cell migration and phagocytosis. Postmortem histological studies of ALSP patients have previously uncovered reduced microglia densities (decreased IBA1+ cells (44–46)) and decreased expression of microglia homeostatic markers including P2RY12 compared to control individuals (44). In contrast, increased number of dysmorphic CD68+ cells have been observed (44, 46). The loss of P2RY12 and increase in CD68 immunoreactivity pointed toward an altered phenotype of resident microglia and/or migration of peripheral monocytes into the ALSP brain. In the current study, lower cell yield upon microglial differentiation of ALSP-CSF1R iPSCs was observed, consistent with the developmental role of CSF1R and histological observations made in ALSP patients. In addition, significantly lower expression of *P2RY12* and higher expression of CD68 in ALSP-CSF1R iMGL was observed compared to controls. This would suggest that phenotypic alterations of microglia, rather than monocyte migration to the brain, are responsible for the histological observations made in ALSP.

High dependency of lineage commitment on CSF1R signaling has been described for microglia and tissue-resident macrophages but not circulating monocytes (8, 47, 48). In the brain, CSF1R is almost exclusively expressed by microglia (49, 50). As such, ALSP is considered a primary microgliopathy. Given the importance of CSF1R signaling throughout microglia development and maintenance, it remains unclear why ALSP typically manifests late in life (43-years-old on average (13)). Aside from CSF1R, another growth factor receptor which promotes microglia proliferation is colony-stimulating factor receptor (CSF2R). Both the CSF1R ligand M-CSF and the CSF2R ligand GM-CSF have been observed to have mitogenic effect on fetal and adult microglia *in vitro* (51). GM-CSF and CSF2R expression is detectable in the human brain during fetal development (52). Yet, whereas *Csf1r* knockout results in an almost complete absence of microglia in mice (8, 12), *Csf2r* knockout does not impact microglia density (53). These observations suggest that CSF1R but not CSF2R is essential for microglia development. GM-CSF is poorly expressed in the adult brain (52, 53), but is upregulated in some inflammatory contexts such as multiple sclerosis (54, 55), in which it is thought to exert its mitogenic effect on microglia. Interestingly, increased GM-CSF expression has been previously detected in the grey matter of ALSP patients (44, 53). This was associated with a less pronounced molecular and morphological alterations of microglia in the grey matter compared to the white matter of those patients (44, 46), suggesting CSF2R signaling might play a compensatory role to CSF1R signaling in ALSP patients. Accordingly, addition of GM-CSF to the microglia differentiation medium was essential for the successful derivation of iMGL from ALSP-CSF1R iPSCs.

CSF1R signaling is known to be important for microglial proliferation in response to stress or injury, with increased M-CSF expression observed upon *in vivo* LPS treatment (56), upon ischemic injury (57), in Alzheimer’s disease and its animal models (56, 58, 59), and in aged mice (56) and human (60). It can be hypothesized that CSF1R-mediated proliferation or functional changes of microglia are of increased importance in maintaining brain homeostasis during aging, explaining the late-onset of ALSP. In mice, aging has been associated with myelin fragmentation and increased sarkosyl-insoluble lipofuscin accumulation in lysosomes of white matter microglia, suggestive of increased lysosomal burden with aging (61). The presence of pigmented lipid-laden or lipofuscin-rich CD68+ myeloid cells in the white matter is a pathological hallmark of ALSP (62–67). Here, ALPS-CSF1R iMGL were observed to have a higher capacity to internalize myelin debris, possibly owing to a higher lysosomal content, compared to control iMGL. Similarly, *Csf1r*^+/-^ microglia have been previously shown to exhibit higher CD68 expression and phagocytosis of myelin compared to microglia from *Csf1r*^+/+^ mice (68). The excess storage of internalized myelin might be responsible for lipid and lipofuscin accumulation within microglia in the ALSP brain.

The association between ALSP and variants located in the coding region of CSF1R’s kinase domain was established in 2011 (39). Given the variable clinical expressivity and incomplete penetrance of *CSF1R* variants observed even within family members sharing the same variant (37, 69, 70), it is likely that other genetic, biological or environmental factors contribute to the severity of the resulting phenotype. Generation of an isogenic control iPSC line in the future will be important in isolating microglial dysfunction caused by *CSF1R* pathogenic variant from that caused by other genetic factors. Generation of iMGL from other ALSP patients will help identify common microglial features specific to ALSP patients. Moreover, unbiased, omic approaches should allow us to better grasp the full picture of phenotypic alterations that are present in ALSP microglia.

The newly devised 2.9 protocol for iMGL generation will be useful not only in the context of ALSP, but also for the *in vitro* study of microglia in any neurological diseases. The new protocol conserved the simplicity of the 2.0 protocol developed by McQuade *et al*., not requiring any cell sorting nor the use of a hypoxia chamber to generate progenitor cells. Composition of the microglia differentiation medium was simplified, requiring less reagents. Generation of iMGL is labor-intensive and involves the use of expensive recombinant proteins. The higher yields achieved by the 2.9 over the 2.0 protocol will allow for a significant cost reduction. In addition, cells differentiated using the 2.9 protocol are more adherent to cultureware, which is convenient when performing image-based assays. Most importantly, transcriptional profile and functional competence of iMGL 2.9 was substantially closer to that of primary microglia, signifying higher translatability of research findings to human disease.

## Conclusions

In conclusion, our newly devised 2.9 protocol for iMGL generation is a promising tool in studying molecular and functional alterations of microglia caused by pathogenic variants associated with human diseases, including primary microgliopathies such as ALSP. This study represents the first successful attempt at investigating iMGL derived from a *CSF1R*-mutated ALSP patient, and revealed molecular alterations that are consistent with previous histopathological findings made in ALSP.

## List of abbreviations

ADP: adenosine diphosphate
ALSP: adult-onset leukoencephalopathy with axonal spheroids and pigmented glia
BSA: bovine serum albumin
CCL3: chemokine ligand 3
CD11b: cluster of differentiation 11b
CD14: cluster of differentiation 14
CD200: cluster of differentiation 200
CD34: cluster of differentiation 34
CD43: cluster of differentiation 43
CD68: cluster of differentiation 68
CSF1R: colony-stimulating factor 1 receptor
CSF2R: colony-stimulating factor 2 receptor
CX3CL1: C-XC-3 motif chemokine ligand 1
CX3CR1: C-XC-3 motif chemokine receptor 1
DMEM: Dulbecco’s Modified Eagle Medium
EDTA: Ethylenediaminetetraacetic acid
FBS: fetal bovine serum
FITC: fluorescein isothiocyanate
GAPDH: glyceraldehyde 3-phosphate dehydrogenase
GAS6: growth arrest specific 6
GM-CSF: granulocyte-macrophage colony-stimulating factor
GPR34: G-protein coupled receptor 34
IBA1: Ionized calcium binding adaptor molecule 1
IFNγ: interferon gamma
iHPC: induced pluripotent stem cell-derived hematopoietic progenitor cell
IL-10: interleukin 10
IL-1Ο: interleukin 1 beta
IL-34: interleukin 34
IL-6: interleukin 6
iMGL: induced pluripotent stem cell-derived microglia
iPSC: induced pluripotent stem cell
LAMP1: lysosomal-associated membrane protein 1
LPS: lipopolysaccharide
LY96: lymphocyte antigen 96
M-CSF: macrophage colony-stimulating factor
MEM: Minimum Essential Medium
MERTK: Mer tyrosine kinase
MMP8: metalloproteinase 8
mRNA: messenger ribonucleic acid
NF-κB: nuclear factor kappa B
NLRP3: NLR family pyrin domain containing 3
OCT-3/4: octamer binding transcription factor 3/4
P2RY12: purinergic receptor P2Y12
PBMC: peripheral blood mononuclear cell
PBMC-Mχπ: peripheral blood mononuclear cell-derived macrophages
PCA: principal component analysis
PROS1: protein S
P/S: penicillin/streptomycin
qPCR: quantitative polymerase chain reaction
qRT-PCR: real-time quantitative reverse transcription polymerase chain reaction
RNA: ribonucleic acid
SDS: sodium dodecyl sulfate
SEM: standard error of the mean
SIGLEC10: sialic acid binding Ig-like lectin 10
SSEA-4: stage-specific embryonic antigen-4
TGF-Ο1: transforming growth factor beta 1
TLR4: toll-like receptor 4
TMEM119: transmembrane protein 119
TNF: tumor necrosis factor
TREM2: triggering receptor expressed on myeloid cell 2
YWHAZ: tyrosine 3-monooxygenase/tryptophan 5-monooxygenase activation protein zeta

## Declarations

### Ethics approval and consent to participate

All tissues or cells were obtained with written consent and under local ethic board’s approval. Use of all human materials was approved by the McGill University Health Centre Research Ethics Board, under project# 1989-178 for human brain tissues and 2019-5374 for iPSCs. Full written informed consent was obtained from the ALSP patient.

### Consent for publication

Not applicable.

### Availability of data and materials

RNAseq data will be made available on Gene Expression Omnibus upon acceptance of this manuscript for publication.

### Authors’ contributions

MFD conceptualized the study and designed all *in vitro* experiments, acquired, analyzed and interpreted data, and prepared the manuscript. DC acquired and analyzed some *in vitro* data, and helped with manuscript preparation in relation to clinical, radiological and genotyping data. MY analyzed RNAseq data. PF acquired and analyzed some *in vitro* data and helped with literature search. CXQC, TMG and VECP were involved in ALSP patient cell reprogramming into iPSCs, expansion and/or quality control procedures. MN carried out and analyzed sequencing data. AM generated PBMC-Mθ>. VECP and JAS contributed to study design. VECP and IS contributed to preliminary testing of ALSP iMGL generation using the 2.0 protocol. RWRD and JAH surgically collected human brain tissues for primary microglia isolation. RLP conceptualized the study, collected and interpreted clinical, radiological and genotyping data, recruited the ALSP patient for iPSC generation and revised the manuscript. LMH and TMD substantively contributed to the design of the study and to critical manuscript revision. All authors read and approved the final version of the manuscript.

## Acknowledgements

We thank lab members Qiao-Ling Cui, Florian Pernin, Genevieve Dorval, Wen Luo, Julien Sirois and Wolfgang Reintsch for technical or administrative assistance.

We thank the patients and their families for their cooperation.

## Additional files

**Additional file 1: Supplementary figures** (Supplementary Figures.pdf) Figures S1-S12.

**Additional file 2: Table S1** (Table S1.xlsx) Information on iPSC sources.

**Additional file 3. Table S2** (Table S2.xlsx)

*CSF1R* variants in the kinase domain coding region detected by Sanger sequencing of iMGL.

