## Additional file 1 for "An adapted stem cell-derived microglia protocol for the study of microgliopathies and other neurological disorders"

**A**

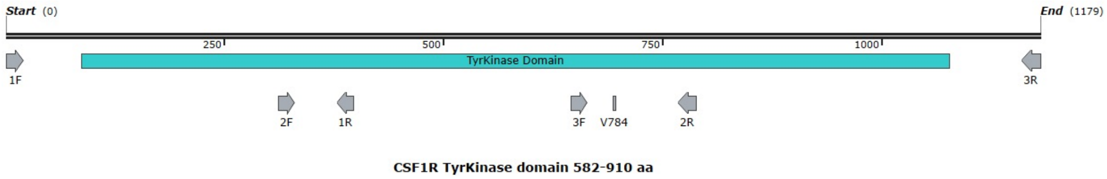


**B**

| Primer name | Sequence (5'-3') |
|---|---|
| CSF1R-TyrKinase-1F | TCGAGAGCTATGAGGGCAAC |
| CSF1R-TyrKinase-1R | GTCCCAGCATGGCCTCAG |
| CSF1R-TyrKinase-2F | AGGCCCTGTACTGGTCATCA |
| CSF1R-TyrKinase-2R | GCATTGCCCTTGACAATGTA |
| CSF1R-TyrKinase-3F | TTCCTCGCTTCCAAGAATTG |
| CSF1R-TyrKinase-3R | TCTCCTCCTCCAGCTCACTG |

**Figure S1. Sequencing strategy of the region encoding CSF1R tyrosine kinase domain.** (A) Locations where the primers would anneal. (B) Sequences of primers.

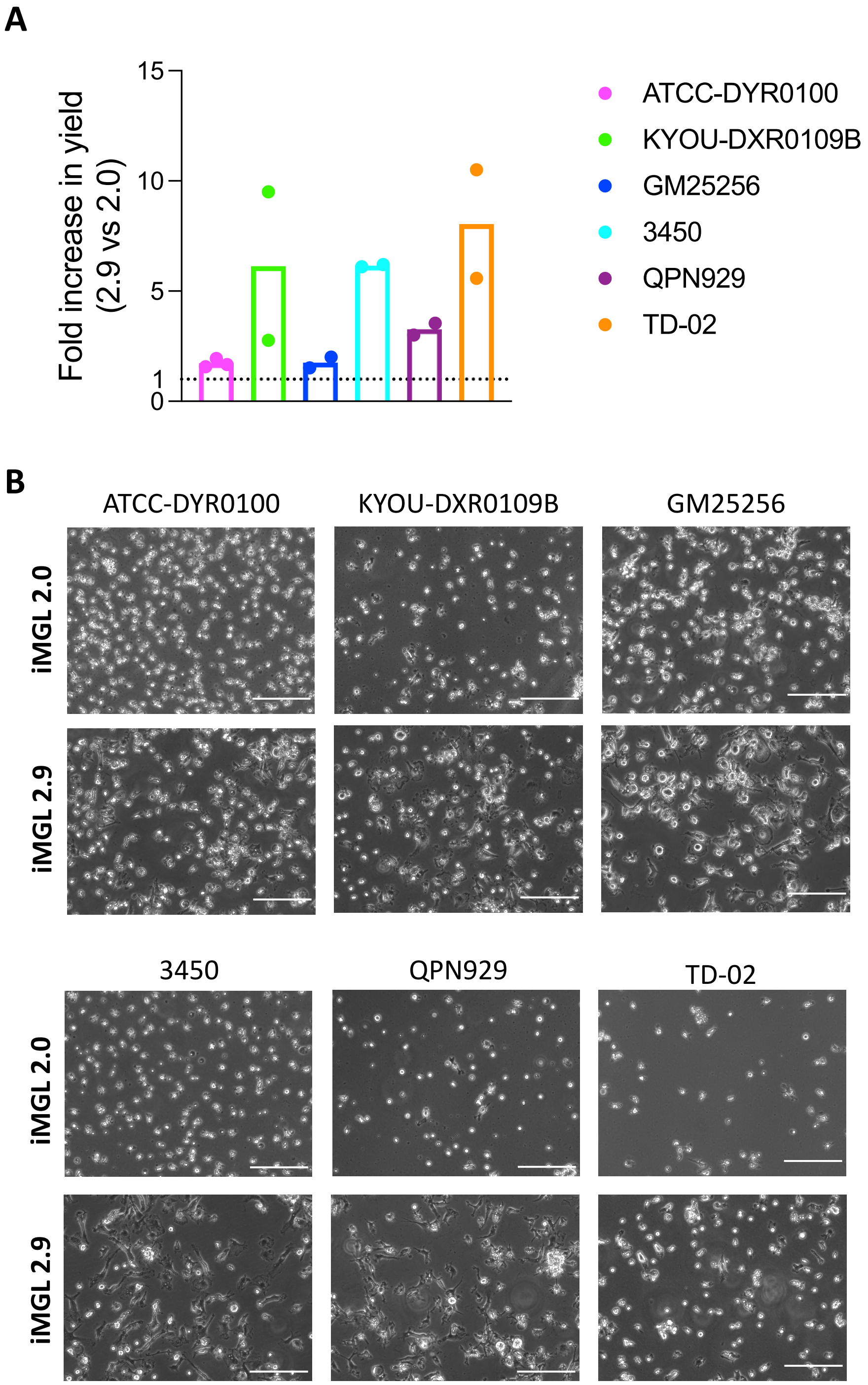


**Figure S2. Cell yield and morphology of iMGL 2.0 and 2.9.** iMGL 2.0 and 2.9 were generated side-by-side from six iPSC lines. (A) Viable cell yield assessed by Trypan blue exclusion assay. n = 2-3 per iPSC line. (B) Phase contrast images. Scale bar = 150 $\mu m$.

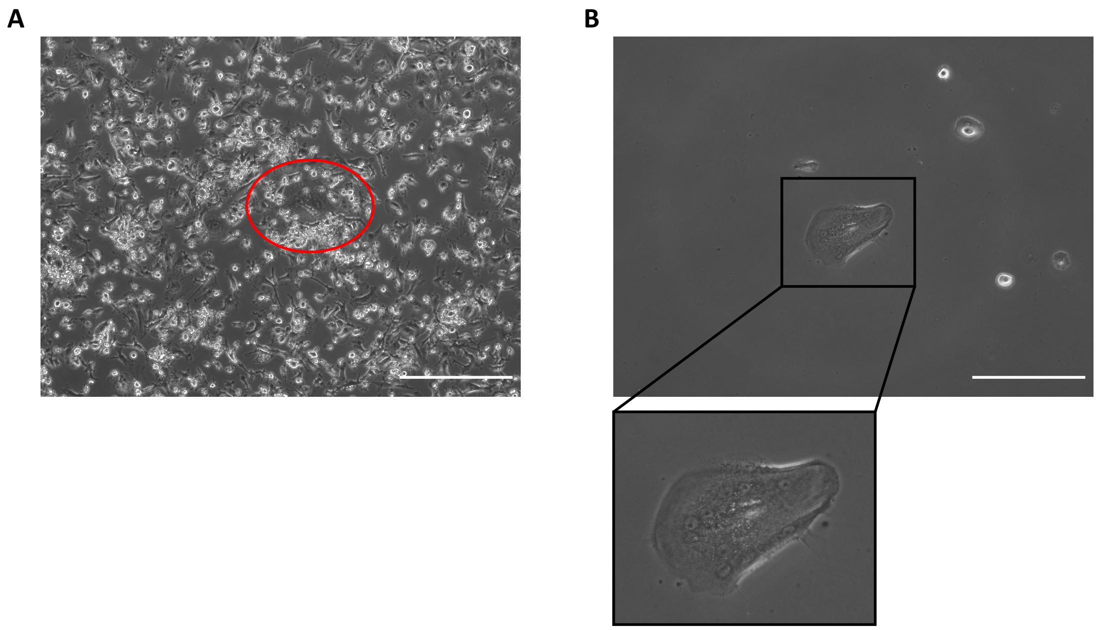


**Figure S3. Multinucleated giant cell contaminants.** Phase contrast pictures of iMGL 2.9 culture before (A) and after (B) EDTA-mediated cell harvest. Red circle in (A) depicts a multinucleated giant cell underneath mononuclear iMGL. Black square in (B) shows a zoomed-in image of a multinucleated giant cell. Scale bar = 300 μm.

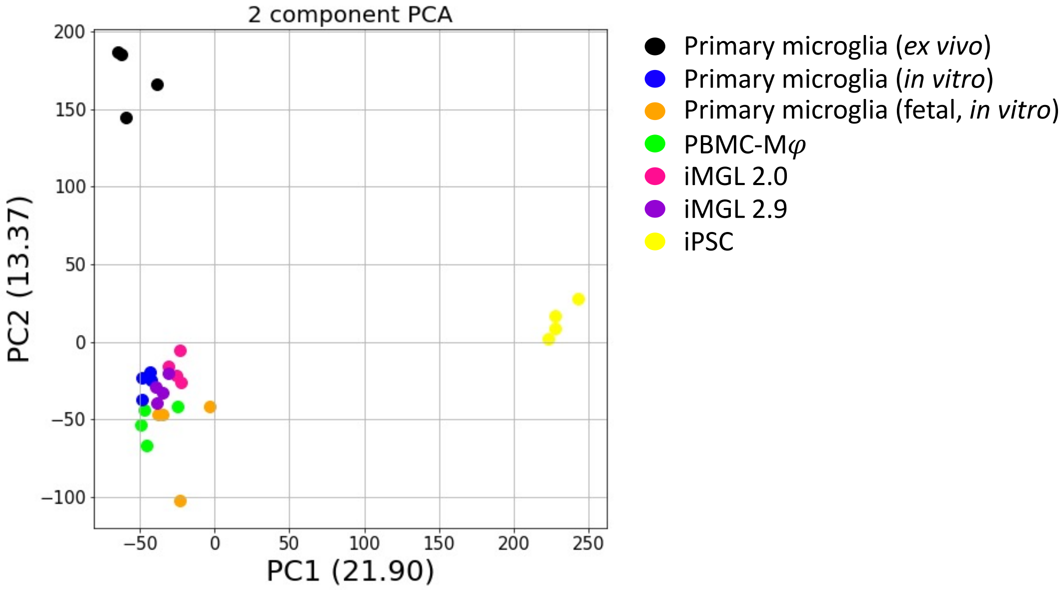


**Figure S4. Transcriptome of iMGL 2.0 and 2.9 compared to primary microglia and iPSCs.** PCA plot of RNA-sequencing data is presented. iMGL 2.0 and 2.9 were differentiated side-by-side from the same iPSC lines.

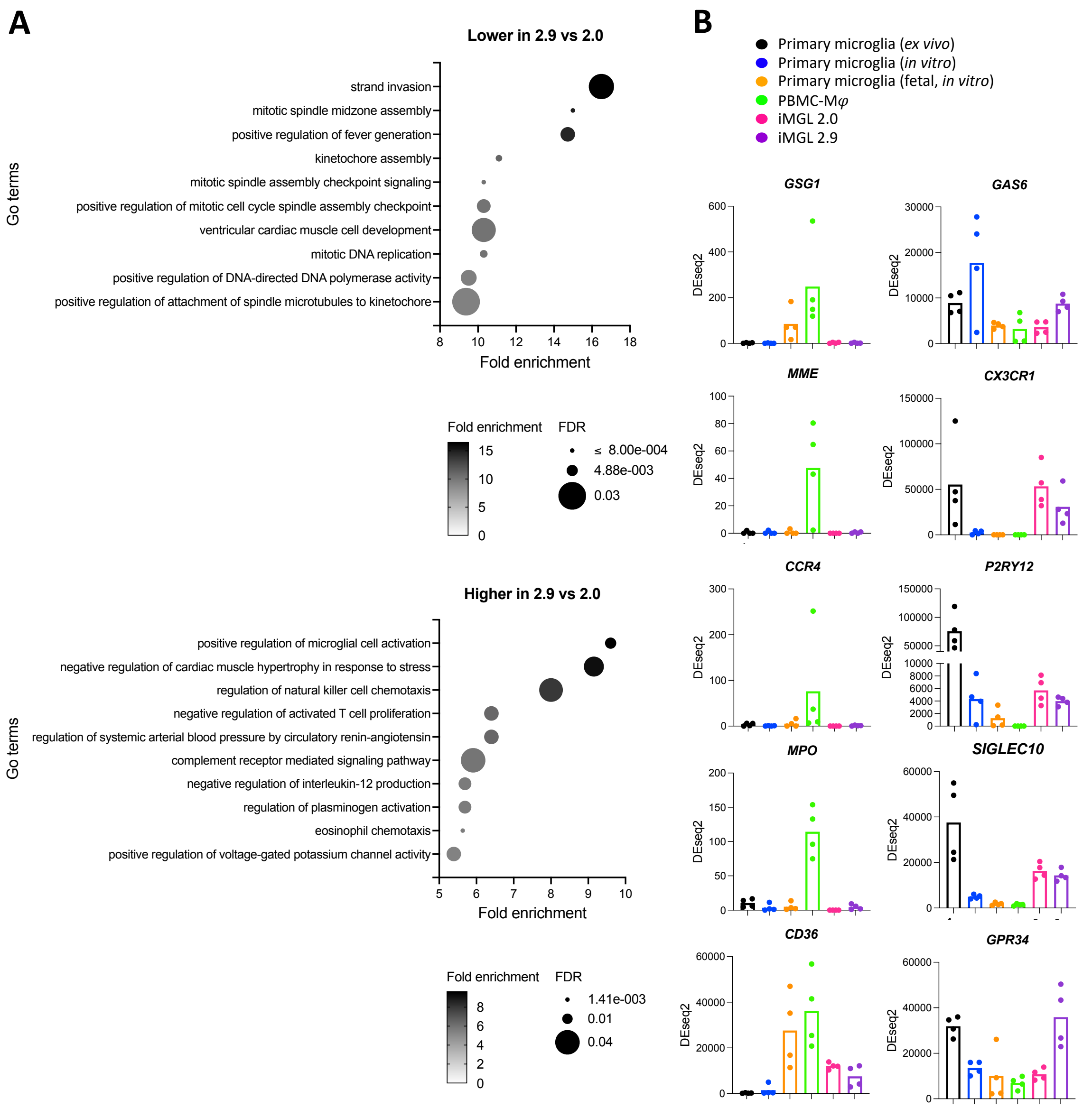


**Figure S5. RNAseq analysis of iMGL 2.9 transcriptome.** iMGL 2.0 and 2.9 were generated side-by-side from the same iPSC lines. (A) GO term analysis of DEGs identified between iMGL 2.9 and 2.0. FDR = false discovery rate. (B) Expression of select macrophage and microglia markers. n = 4 donors for primary cells, 4 differentiation batches from 4 iPSC lines for iMGL.

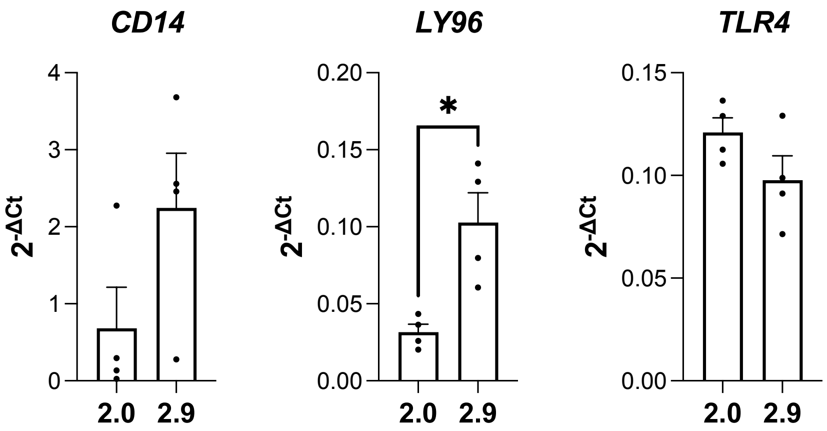


**Figure S6. qRT-PCR assessment of *CD14*, *LY96* and *TLR4* expression in iMGL 2.0 and 2.9.** iMGL 2.0 and 2.9 were differentiated side-by-side from the same iPSC lines. T-tests were performed. n = 4 differentiation batches from 4 iPSC lines, * $p < 0.05$.

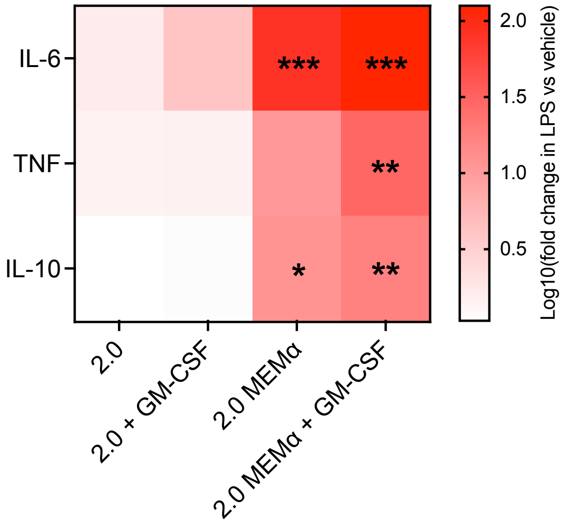


**Figure S7. Media-dependent inflammatory response of iMGL to LPS.** iMGL were differentiated using the indicated media formulation and then treated with vehicle or LPS (100 ng/mL) for 24 hours. Cytokine secretion in cell supernatants was assessed. A two-way ANOVA was performed, followed by Dunnett's post hoc test. n = 3 differentiation batches from 3 different lines, * $p < 0.05$, ** $p < 0.01$, *** $p < 0.001$ versus iMGL 2.0.

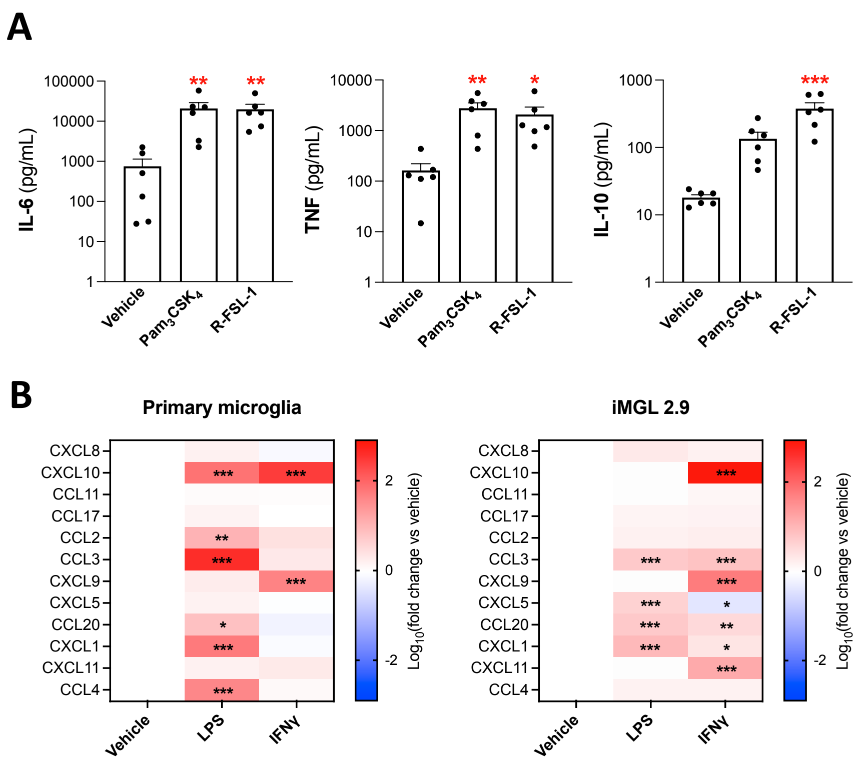


**Figure S8. Inflammatory response of iMGL 2.9 to a variety of stimuli.** (A) IL-6, TNF and IL-10 concentrations in supernatants from iMGL 2.9 treated with vehicle, $Pam_3CSK_4$ (100 ng/mL) or R-FSL-1 (250 ng/mL) for 24 hours. One-way ANOVA were performed, followed by Dunnett's post hoc tests. n = 4 differentiation batches from 6 iPSC lines, * $p < 0.05$, ** $p < 0.01$ versus vehicle. (B) Heatmap representing chemokine secretion from primary human microglia or iMGL 2.9 following a 24-hour treatment with LPS (100 ng/mL) or IFNγ (10 ng/mL). Two-way ANOVA were performed, followed by Dunnett's post hoc tests. n = 2 donors for primary microglia and 3 differentiation batches from 3 iPSC lines for iMGL. * $p < 0.05$, ** $p < 0.01$, *** $p < 0.001$ versus vehicle.

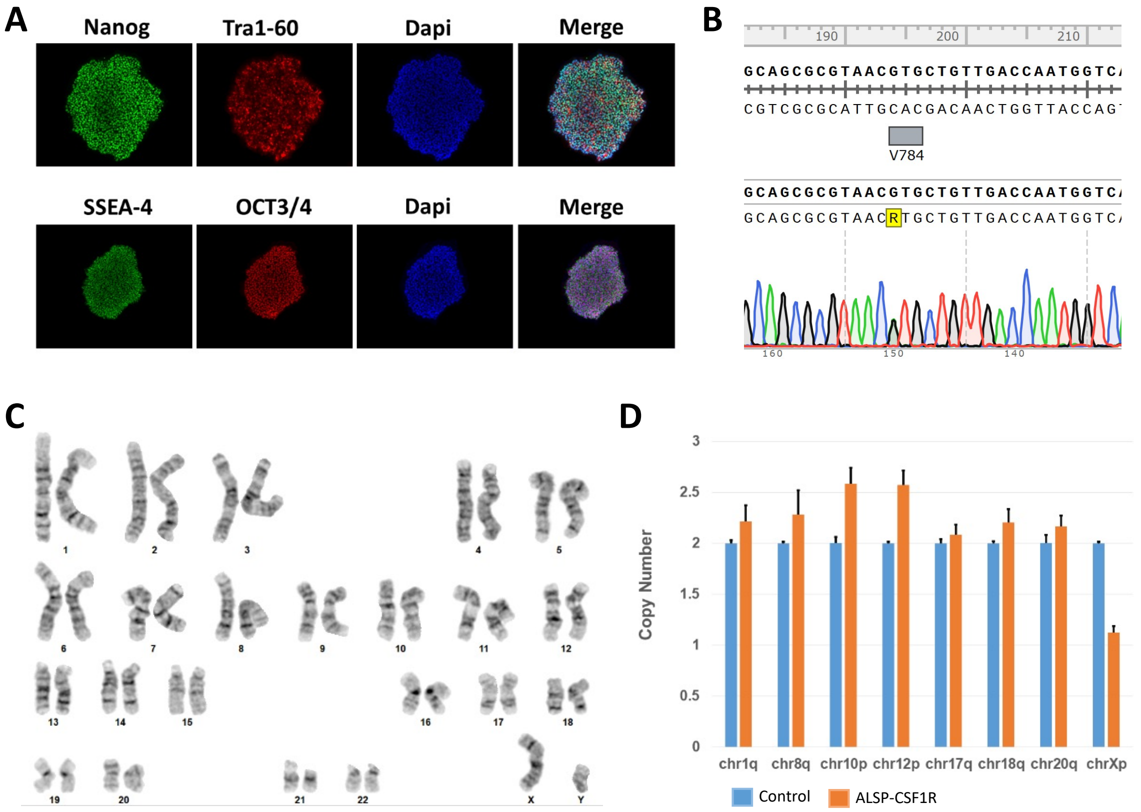


**Figure S9. Quality control of the ALSP-CSF1R iPSC line.** (A) Immunostaining of pluripotency markers. (B) Chromatogram of the heterozygous c.2350G > A variant in the *CSF1R* gene (shown here at position 150). (C) G-band assay showing normal karyotype of ALSP-CSF1R iPSCs. (D) qPCR-based assessment of copy numbers of chromosomal regions prone to karyotypic abnormalities.

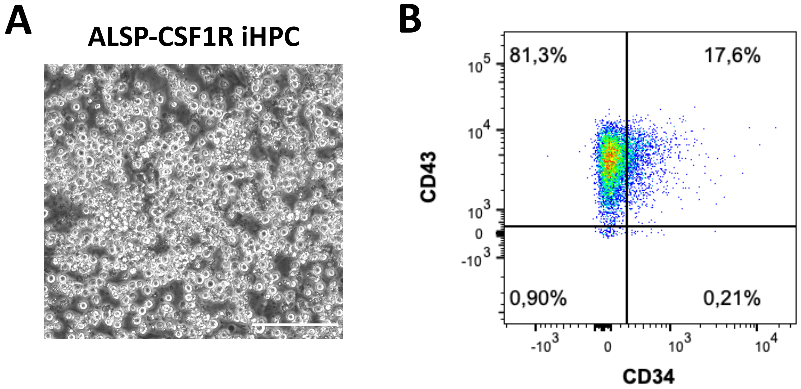


**Figure S10. iHPCs generated from ALSP-CSF1R iPSCs.** (A) Phase contrast image of iHPC culture on day 12 of differentiation (scale bar = 150 μm). (B) Flow cytometry assessment of CD43 and CD34 in iHPCs on day 12 of differentiation.

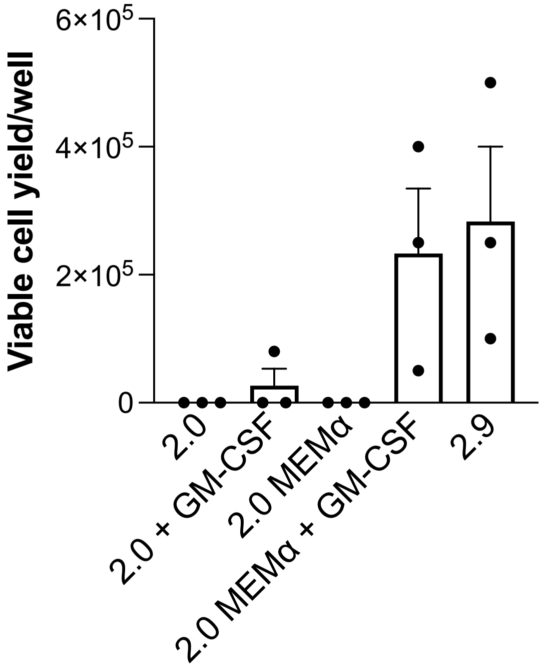


**Figure S11. Cell yield following microglial differentiation of ALSP-CSF1R cells using various media formulations.** Viable cell yield per well of a 6-well plate assessed by trypan blue exclusion assay. n = 3 differentiation batches.

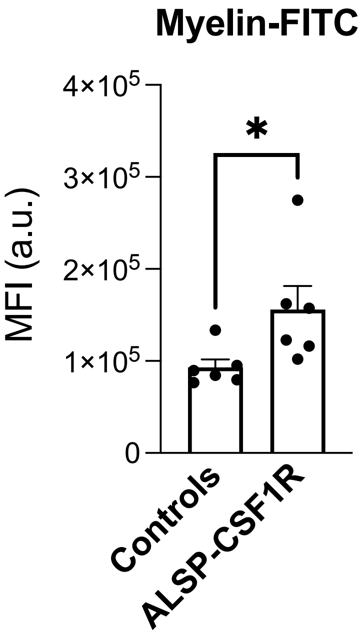


**Figure S12. Internalization of FITC-labelled myelin by ALSP-CSF1R iMGL compared to control iMGL.** iMGL 2.9 were exposed to FITC-labelled myelin for three hours and then extracellular fluorescence was quenched by trypan blue. Cells were counterstained with Hoechst 33342 and mean green fluorescence intensity (MFI) was measured (a.u. = arbitrary unit). A t-test was performed. n = 6 batches from 4 healthy control iPSC lines and 6 batches of a single ALSP patient line, differentiated side-by-side, * $p < 0.05$.
